# Repurposing the antidepressant sertraline as SHMT inhibitor to suppress serine/glycine synthesis addicted breast tumor growth

**DOI:** 10.1101/2020.06.12.148395

**Authors:** Shauni L. Geeraerts, Kim R. Kampen, Gianmarco Rinaldi, Purvi Gupta, Mélanie Planque, Kaat De Cremer, Katrijn De Brucker, Stijn Vereecke, Benno Verbelen, Pieter Vermeersch, David Cassiman, Sarah-Maria Fendt, Arnout Voet, Bruno P.A. Cammue, Karin Thevissen, Kim De Keersmaecker

## Abstract

Metabolic rewiring is a hallmark of cancer that supports tumor growth, survival and chemotherapy resistance. While normal cells often rely on extracellular serine and glycine supply, a significant subset of cancers becomes addicted to intracellular serine/glycine synthesis, offering an attractive drug target. Previously developed inhibitors of serine/glycine synthesis enzymes did not reach clinical trials due to unfavorable pharmacokinetic profiles, implying that further efforts to identify clinically applicable drugs targeting this pathway are required. In this study, we aimed to develop therapies that can rapidly enter the clinical practice by focusing on drug repurposing, as their safety and cost-effectiveness have been optimized before. Using a yeast model system, we repurposed two compounds, sertraline and thimerosal, for their selective toxicity against serine/glycine synthesis addicted breast cancer and T-cell acute lymphoblastic leukemia cell lines. Isotope tracer metabolomics, computational docking studies and an enzymatic activity assay revealed that sertraline and thimerosal inhibit serine/glycine synthesis enzymes serine hydroxymethyltransferase and phosphoglycerate dehydrogenase, respectively. In addition, we demonstrated that sertraline’s anti-proliferative activity was further aggravated by mitochondrial inhibitors, such as the antimalarial artemether, by causing G1-S cell cycle arrest. Most notably, this combination also resulted in serine-selective antitumor activity in breast cancer mouse xenografts. Collectively, this study provides molecular insights into the repurposed mode-of-action of the antidepressant sertraline and allows to delineate a hitherto unidentified group of cancers being particularly sensitive to treatment with sertraline. Furthermore, we highlight the simultaneous inhibition of serine/glycine synthesis and mitochondrial metabolism as a novel treatment strategy for serine/glycine synthesis addicted cancers.

## INTRODUCTION

Rewiring of energy metabolism, exemplified by the well-established Warburg effect, is one of the ten hallmarks of cancer (1). While normal cells often rely on serine and glycine uptake from their environment, several cancer subtypes produce their own serine and glycine via intracellular serine/glycine synthesis and become addicted to this own production (2,3). In general, serine/glycine synthesis (Figure 1) consists of two processes: *de novo* serine synthesis from glucose and interconversion of serine into glycine. *De novo* serine synthesis branches from glycolysis, with the glycolytic intermediate 3-phosphoglycerate being converted into serine via three consecutive enzymatic reactions catalyzed by phosphoglycerate dehydrogenase (PHGDH), phosphoserine aminotransferase 1 (PSAT1) and phosphoserine phosphatase (PSPH). Thereafter, serine is catabolized into glycine, facilitated by serine hydroxymethyltransferase 1 or 2 (SHMT1/2), with 1 and 2 referring to the cell compartment in which the reaction takes place, i.e. the cytosol or the mitochondria (2,3). By relying on intracellular serine/glycine synthesis, cancer cells feed their high requirements to generate formate for purine synthesis, produce reductive equivalents to control redox homeostasis, regulate DNA demethylation and support lipid metabolism (3).

**Figure 1.**
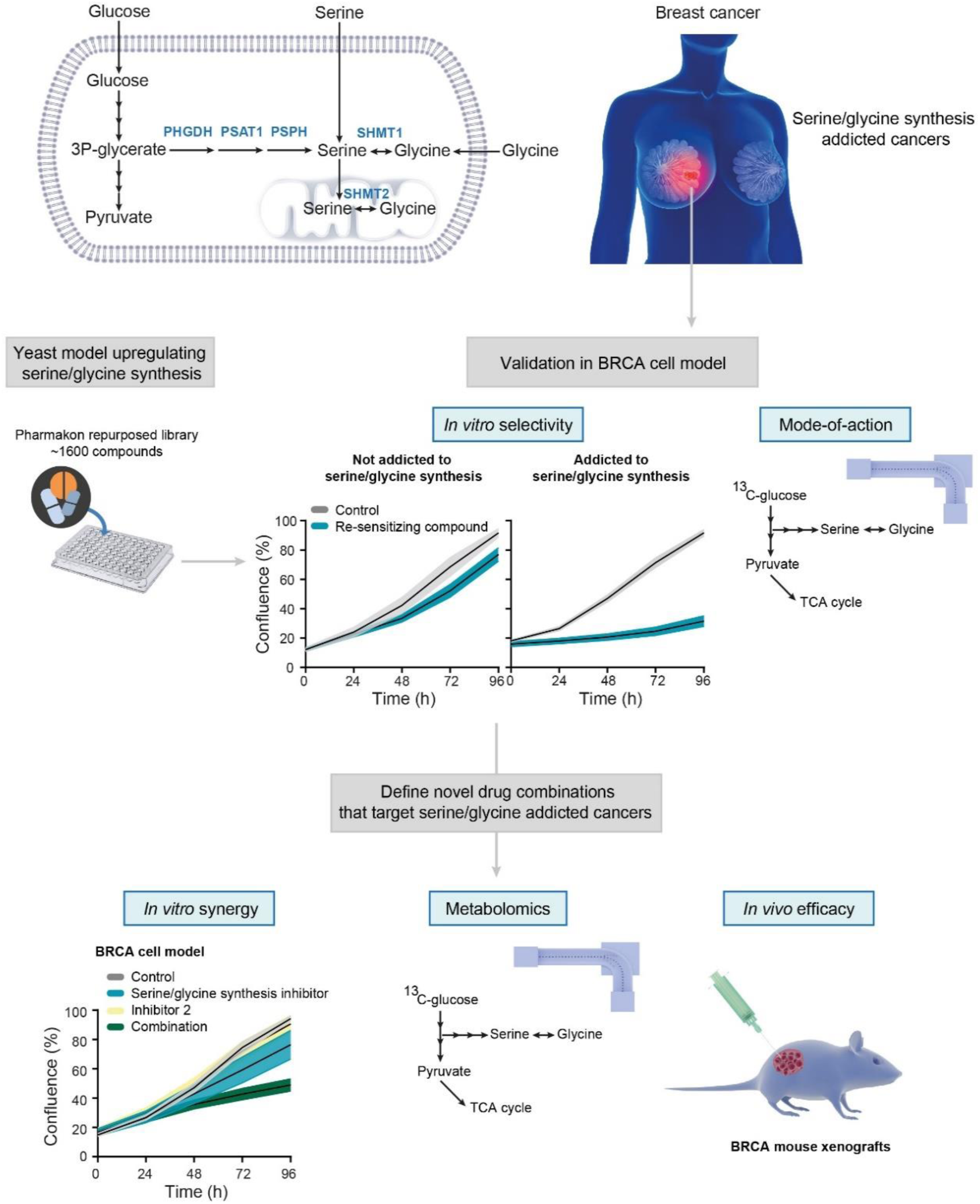
Schematic overview of study design. In terms of serine/glycine metabolism, breast cancer (BRCA) can largely be divided into serine/glycine uptake or synthesis addicted. Using a lower eukaryotic yeast model system that upregulates serine/glycine synthesis, we selected repurposed compounds that target serine/glycine synthesis in the breast cancer context. After thorough *in vitro* validation, the most promising, and clinically used, repurposed compound was selected for the rational design of a novel combination therapy for serine/glycine synthesis addicted breast cancer. Human enzymes involved in serine/glycine synthesis are indicated in blue. PHGDH: phosphoglycerate dehydrogenase; PSAT1: phosphoserine aminotransferase; PSPH: phosphoserine phosphatase; SHMT1/2: cytosolic/mitochondrial serine hydroxymethyltransferase.

A cancer type well-known for its dependency on serine/glycine synthesis is breast cancer, where 6% of the patient samples display copy number gains of the *PHGDH* gene. Furthermore, 70% of estrogen receptor negative breast tumors have increased PHGDH protein levels, and inhibition of PHGDH via RNA interference or PHGDH inhibitors impairs cell proliferation and survival (4–8). Besides PHGDH, also SHMT2 is known to be exploited by breast cancer cells as SHMT2 expression levels are positively correlated with breast cancer grade (9). Additionally, SHMT2 is identified as a direct target gene of the MYC oncogene (10). MYC-driven stimulation of *de novo* serine synthesis by transcriptional upregulation of PHGDH, PSAT1 and PSPH is also critical for sustaining survival and rapid proliferation of cancer cells under nutrient deprived conditions (11). Apart from MYC, the oncogene KRAS, the tumor suppressor p53, the mTOR-ATF4 axis and the T-cell leukemia associated R98S mutation in ribosomal protein L10 (RPL10 R98S) have also been shown to enhance serine/glycine synthesis in cancer cells (12–15).

Evidence for dependency on serine/glycine synthesis in cancer subsets, including triple-negative breast cancer and T-cell leukemia that are both currently treated with toxic intensive chemotherapy regimens, is growing. This highlights the necessity to develop novel therapeutic intervention strategies for these cancers, specifically focusing on targeting serine/glycine synthesis. PHGDH and SHMT inhibitors have been identified but did not enter clinical trials due to unfavorable pharmacokinetic profiles or because they have only recently been developed (7,8,16,17). Consequently, this addresses the need for clinically applicable drugs targeting this pathway.

In this study, we made use of a lower eukaryotic yeast model system that specifically upregulates serine/glycine synthesis in response to sublethal stress. Using this platform, we discovered two repurposed compounds, sertraline and thimerosal, that show selective toxicity to serine/glycine synthesis addicted cancer cell lines, while they had no effect on cancer and normal lymphoid cell lines that take up serine and glycine from their environment. Moreover, this work reports the potential application of the widely clinically used antidepressant sertraline as adjuvant therapeutic agent to treat serine/glycine synthesis addicted cancers, especially when combined with drugs causing general mitochondrial dysfunction.

## MATERIALS AND METHODS

### Human cell cultures

Breast cancer cell lines MDA-MB-231, MDA-MB-468, MCF7 and HCC70 (American Type Culture Collection; ATCC) were cultured in DMEM medium (Life Technologies) supplemented with 10% fetal bovine serum (FBS; Life Technologies). Because of many serial passages, MCF7 cell authenticity was confirmed by Microsynth AG. Other cell lines were freshly obtained from ATCC. The Ba/F3 pro-B cell line was obtained from Leibniz-Institute DSMZ and grown in RPMI-1640 (Life Technologies) with 10% FBS (Life technologies). All cells were mycoplasma negative. Serine depleted media has the following composition: DMEM without serine and glycine (US Biological life Sciences, D9802-01), supplemented with 4.5 g/l glucose (Sigma-Aldrich), 3.7 g/l sodium bicarbonate (Sigma-Aldrich), glutamax (100x, Life Technologies) and 10% dialyzed serum (Life Technologies, A3382001).

### Compounds

1000x stock solutions of each compound were made and stored at -20 °C. Sertraline (Sigma-Aldrich), NCT-503 (Sigma-Aldrich), rotenone (Sanbio), antimycin A (Sigma-Aldrich) and artemether (TCI Europe) were dissolved in DMSO (Sigma-Aldrich), while thimerosal (Sigma-Aldrich), benzalkonium chloride (Sigma-Aldrich) and bupropion (TCI Europe) were dissolved in water (Invitrogen). Sertraline’s activity can differ between batches, causing variability in the exact dose that is required to obtain the observed effects.

### Cancer cell proliferation assay

MDA-MB-231 (7500 cells/well), MDA-MB-468 (7500 cells/well), MCF7 (4000 cells/well) and HCC70 (4500 cells/well) breast cancer cells were plated in 100 µl of DMEM culture medium (Life Technologies) in 96-well plates (TPP 96 well tissue culture plates from Sigma-Aldrich) and incubated for 24 h at 37 °C to obtain optimal adherence to the surface. Then, 100 µl of 2x solutions of the compound solutions, diluted in DMEM culture medium with 10% FBS, were added to the cells. During the following 5 days, cell proliferation was assessed by real-time live-cell imaging analysis of confluency on an IncuCyte Zoom system (Essen BioScience). Using the collected data, we calculated the growth rate under various drug conditions. Only the exponential (logarithmic) portions of the resulting growth curves were used for determining the growth rates. To do this, the formula below was used, in which T1 and T2 are two time points within the exponential growth phase.

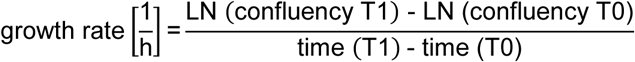

### Generation of Ba/F3 CRISPR-Cas9 RPL10 R98S mutant cells

Ba/F3 cells (ATCC) were electroporated (6 square wave pulses, 0.1ms interval, 175V) with a pX458 vector (a gift from Feng Zhang; Addgene plasmid #48138) containing an Rpl10 targeting gRNA (5’-TGATACGGATGACATGGAAA-3’) and a 117-nt donor oligo containing the R98S allele as well as 3 silent mutations to avoid re-recognition and cutting by the gRNA-Cas9 complex (5’-CCAACAAATACATGGTAAAGAGTTGTGGCAAGGATGGCTTTCATATCCGAGTGAGGCTCCATCCTTTTCATGTAATCAGTATCAACAAGATGTTGTCCTGTGCTGGGGCTGACAGGT-3’; Integrated DNA Technologies). Following electroporation, cells were incubated for 24 h in the presence of 10 µM SCR7 (Sigma-Aldrich), followed by single cell sorting (BD FACS Aria III) into ClonaCell™-TCS Medium (STEMCELL Technologies). Outgrowing clones were expanded and screened for the desired modification using Sanger sequencing to confirm the Rpl10 status and to check for modification of relevant predicted off-target effects (Rpl10-ps3 and Rpl10l).

### Ba/F3 RPL10 WT and R98S viability assay

Three RPL10 WT versus three RPL10 R98S clones were plated and cultured for 72 h to exhaust the medium. After these 72 h, the clones were incubated with sertraline (7.3 µM) and thimerosal (1 µM). Relative cell viability was measured, using flow cytometry, after 48 h treatment with the compounds.

### Steady-state metabolite concentrations and _13_C_6_-glucose tracing

A number of 150.000 MDA-MB-468 cells were plated in 3 ml of DMEM culture medium (Life Technologies) in 6-well plates (Greiner Bio-One). After 24 h of incubation at 37 °C, cells were washed with PBS and 3 ml of tracing medium (glucose-free DMEM medium supplemented with 10% dialyzed serum (Life Technologies) and 4.5 g/l ^13^C_6_-glucose (Sigma-Aldrich)) was added. Subsequently, cells were incubated during 24 h (thimerosal and NCT-503) or 72 h (sertraline, artemether and combination) at 37 °C. Metabolites for the subsequent mass spectrometry analysis were prepared by quenching the cells in liquid nitrogen followed by a cold two-phase methanol-water-chloroform extraction (18,19). Phase separation was achieved by centrifugation at 4 °C (24 × 3.75 g, 10 min). The methanol-water phase containing polar metabolites was separated and dried using a vacuum concentrator at 4 °C overnight. Dried metabolite samples were stored at −80 °C. Polar metabolites were analyzed by GC-MS and LC-MS.

### Computational docking studies

Sertraline was modelled using MOE (Chemical Computing Group, Montreal, Canada) with the MMFF94x force field. The structures of the putative receptors present in serine/glycine synthesis were obtained from the RCSB database (www.rcsb.org) (PDB ID SHMT1: 1BJ4, SHMT2: 5V7I). The bioactive conformations were chosen for each receptor (SHMT1, SHMT2 as dimers) and optimized in MOE using protonate_3D. Docking was performed using GOLD8 (20) software. Specifically, sertraline was docked in the presence of the pyrodoxal 5’-phosphate (PLP) co-factor. Furthermore, another known inhibitor of SHMT (pyrazolopyran scaffold) was also docked in the presence of the PLP ligand. Conformational restraints were applied to the ligand by disallowing the flipping of ring conformations and planar R-NR1R2 groups to ensure the rigidity of sertraline. The ligand was docked 10 times into each receptor and the score was calculated using CHEMPLP scoring function. The interactions of sertraline with each receptor were analyzed and visualized using PYMOL version 1.8 (Schrödinger, 2015). The docked conformations of the ligands in SHMT1 and SHMT2 were superimposed using PYMOL to compare the interactions.

### PHGDH *in vitro* enzymatic activity assay

PHGDH enzyme activity upon drug treatment was tested using human PHGDH (BPS bioscience, 71079) and a specific colorimetric PHGDH activity kit (Biovision, K569). The known PHGDH inhibitor, NCT-503, served as a positive control (7). Human PHGDH enzyme was diluted in water (Invitrogen) at a concentration of 0.15 mg/ml. Next, 5 µl of this PHGDH enzyme solution and 10 µl of sertraline, thimerosal or NCT-503 was added in a 96-well plate (flat bottom). Subsequently, 35 µl PHGDH assay buffer (Biovision) and 50 µl PHGDH reaction mix (prepared as described in the protocol of the PHGDH assay kit K569, Biovision) was added. Afterwards, absorbance at 450 nm was measured over time. In between measurements, the plate was incubated at 37 °C, protected from light. Finally, PHGDH activity was calculated between two time point within the linear range of the internal standard (as described in the protocol of the PHGDH assay kit K569, Biovision).

### Deuterated [2,3,3-^2^H]-serine tracing

A number of 150.000 MDA-MB-468 cells were plated in 2 ml of DMEM culture medium (Life Technologies) in 6-well plates (Greiner Bio-One). After incubation for 24 h at 37 °C, cells were washed with PBS and 2 ml of serine-free DMEM (US Biological life Sciences, D9802-01), supplemented with 4.5 g/l glucose (Sigma-Aldrich), 3.7 g/l sodium bicarbonate (Sigma-Aldrich), 400 µM [2,3,3-^2^H]-serine (deuterium labeled serine, Sigma-Aldrich), 400 µM glycine (Sigma-Aldrich), glutamax (100x, Life Technologies) and 10% dialyzed serum (Life Technologies) was added for 48 h. Metabolites for the subsequent mass spectrometry analysis were prepared by quenching the cells in liquid nitrogen followed by a cold two-phase methanol-water-chloroform extraction (18,19). Phase separation was achieved by centrifugation at 4 °C (24 × 3.75 g, 10 min). The methanol-water phase containing polar metabolites was separated and dried using a vacuum concentrator at 4 °C overnight. Dried metabolite samples were stored at −80 °C. Polar metabolites were analyzed by GC-MS and LC-MS.

### Uptake and secretion rates

A number of 150.000 MDA-MB-468 cells were plated in 2 ml of DMEM culture medium (Life Technologies) in 6-well plates (Greiner Bio-One). The day after (day 0), cells were washed with PBS and 2 ml of fresh DMEM medium was added. 72 h later (day 3), medium samples were taken (0.5-1 ml). The cells were counted on day 0 and after 72 h (day 3), using an automated cell counter. Medium samples were analyzed by mass spectrometry. Specifically, uptake and secretion rates were calculated by subtracting metabolite concentrations of treated samples from metabolite concentrations of culture media incubated for the same amount of time but without cells. Finally, results were normalized for both cell number and growth rate.

### Gas chromatography–mass spectrometry (GC-MS)

Polar metabolites were derivatized and measured as described before (18,19). In brief, polar metabolites were derivatized with 20 mg/ml methoxyamine in pyridine for 90 min at 37 °C and subsequently with N-(tert-butyldimethylsilyl)-N-methyl-trifluoroacetamide, with 1% tert-butyldimethylchlorosilane for 60 min at 60 °C. Metabolites were measured with a 7890A GC system (Agilent Technologies) combined with a 5975C Inert MS system (Agilent Technologies). One microliter of samples was injected in split mode (ratio 1 to 3) with an inlet temperature of 270 °C onto a DB35MS column. The carrier gas was helium with a flow rate of 1 ml/min. For the measurement of polar metabolites from the deuterated serine labeling experiment, the GC oven was set at 100 °C for 1 min and then increased to 105 °C at 2.5 °C/min and with a gradient of 2.5 °C/ min finally to 320 °C at 22 °C/min. The measurement of metabolites has been performed under electron impact ionization at 70 eV using a selected-ion monitoring (SIM) mode. For the measurement of polar metabolites from the ^13^C_6_-glucose labeling experiment, the GC oven was held at 100 °C for 3 min and then ramped to 300 °C with a gradient of 2.5 °C/min. The mass spectrometer system was operated under electron impact ionization at 70 eV and a mass range of 100–650 a.m.u. was scanned. Mass distribution vectors were extracted from the raw ion chromatograms using a custom Matlab M-file, which applies consistent integration bounds and baseline correction to each ion (21). Moreover, data were corrected for naturally occurring isotopes (22). For metabolite levels, arbitrary units of the metabolites of interest were normalized to glutaric acid, an internal standard, and protein content.

### Liquid chromatography–mass spectrometry (LC-MS)

A liquid chromatography, Dionex UltiMate 3000 LC System (Thermo Scientific), coupled to mass spectrometry (MS), a Q Exactive Orbitrap (Thermo Scientific), was used for the separation of polar metabolites. A volume of 15 μl of sample was injected and metabolites were separated on a C18 column (Acquity UPLC HSS T3 1.8 µm 2.1×100 mm) at a flow rate of 0.25 ml/min, at 40 °C. A gradient was applied for 40 min (solvent A: 0 H2O, 10 mM Tributyl-Amine, 15 mM acetic acid – solvent B: Methanol) to separate the targeted metabolites (0 min: 0% B, 2 min: 0% B, 7 min: 37% B, 14 min: 41% B, 26 min: 100% B, 30 min: 100% B, 31 min: 0% B; 40 min: 0% B). The MS operated in full scan in negative mode (m/z range: 70-1050 and 300-800 from 8 to 25 min) using a spray voltage of 4.9 kV, capillary temperature of 320 °C, sheath gas at 50.0, auxiliary gas at 10.0. Data was collected using the Xcalibur software (Thermo Scientific) and analyzed with Matlab using the same procedure as described above for the analysis of GC-MS data.

### Flow cytometry – Cell death analysis

Effects on cell death were quantified using a combined Zombie Aqua – eFluor506 and Annexin V – PE staining. A number of 150.000 MDA-MB-468 cells were plated in 2 ml of DMEM culture medium (Life Technologies) in 6-well plates (Greiner Bio-One) and incubated for 24 h at 37 °C to obtain optimal adherence to the surface. The day after, cells were treated with 2 ml of fresh DMEM medium containing sertraline (5, 7.5 or 10 µM) and/or artemether (80 µM). Upon 24 h of treatment, cells were collected in a 96-well U-bottom plate and washed with PBS. Afterwards, cell viability was analyzed following incubation with Zombie Aqua (BioLegend #423102, 1:1000 in PBS) for 20 min at RT in the dark, an Annexin V binding buffer (Thermo Fisher Scientific) washing step and Annexin V (IQ Products IQP-120R, 1:100 in binding buffer) staining for 15 min at RT in the dark. All samples were acquired using a FACS Canto II flow cytometer and analyzed with FlowJo V10 software.

### Flow cytometry – Cell cycle analysis

In general, cell cycle was analyzed by bromodeoxyuridine (BrdU) incorporation and propidium iodide (PI) staining. A number of 150.000 MDA-MB-468 cells were plated in 2 ml of DMEM culture medium (Life Technologies) in 6-well plates (Greiner Bio-One) and incubated for 24 h at 37 °C to obtain optimal adherence to the surface. The day after, cells were treated with 2 ml of fresh DMEM medium containing sertraline (5, 7.5 or 10 µM) and/or artemether (80 µM). Upon 24 h of treatment, cells were incubated with 10 µM BrdU (eBioscience kit #00-4440-51A) for 1 h at 37 °C. Next, cells were collected using trypsin and washed with PBS. Afterwards, BrdU incorporation and PI staining were analyzed following 70% ice cold ethanol fixation, 2 M HCL denaturation, 0.5 M EDTA (pH 8.0) neutralization, BrdU-FITC antibody staining for 20 min at RT (BioLegend #364104) and 20 min incubation with PI solution (100 µg/ml) at RT. All samples were acquired using a FACS Canto II flow cytometer and analyzed with FlowJo V10 software.

### Xenografts in NOD-SCID/IL2γ-/- (NSG) mice

Animal experiments were approved by the local ethics committee (P262-2015). 3×10^6^ breast cancer cells were injected subcutaneously in the left (MDA-MB-231) and right (MDA-MB-468) flanks of NOD-SCID/IL2γ-/- (NSG) mice in a 1:1 mixture with matrigel (Corning). The animals were monitored on a daily basis and sacrificed after 28 days. Mice received treatments on days 7, 9, 11, 13, 15, 20 and 24. Therapy was administered via intra-peritoneal injections at dosages of 2.5 mg/kg sertraline (Sigma-Aldrich) and/or 40 mg/kg artemether (TCI Europe). Control mice were treated with DMSO.

### Statistics

All statistical analyses were performed using GraphPad Prism 8 software and data are presented as mean ± standard deviation (SD). Specific statistical tests used for each experiment are mentioned in the figure legends. Results were considered to be statistically significant if the adjusted p-value was < 0.05 (*p < 0.05, **p < 0.01, ***p < 0.001, ****p < 0.0001).

## RESULTS

### Identification of selective inhibitors of serine/glycine synthesis addicted breast cancer cell lines

Because of the increasing number of identified cancer subsets that are addicted to serine/glycine synthesis, inhibitors that target this pathway are of great interest. Previously identified PHGDH and SHMT inhibitors did not reach clinical trials since they were not able to efficiently target serine/glycine synthesis *in vivo* or because they have only recently been developed (7,8,16,17,23). Therefore, further efforts to identify clinically applicable drugs targeting this pathway are required. To address this need, we focused on drug repurposing with the aim to develop novel therapies that can rapidly enter the clinical practice as their safety and pharmacokinetics have been validated in patients before. To do this, we made use of the lower eukaryotic yeast *Candida albicans* that specifically upregulates serine/glycine synthesis as a tolerance mechanism against sublethal antifungal stress (Figure S1A-B) (24,25), and hypothesized that compounds re-sensitizing this yeast to the antifungal stress, can be potent inhibitors of serine/glycine synthesis. Subsequently, “re-sensitizing hits” were further validated in a cancer context, both *in vitro* and *in vivo*, by investigating their selective anti-proliferative effects on serine/glycine synthesis addicted cancer cell lines and delineating their exact mode-of-action on the serine/glycine synthesis pathway (Figure 1). Interestingly, the serotonin reuptake inhibitor sertraline was identified as a re-sensitizing agent in this lower eukaryotic model system, implying that it might stem from inhibition of serine/glycine synthesis (Figure S1C). This drug caught our attention, as sertraline is widely used in the clinic as antidepressant of which its safety for chronic use is well-documented.

To further investigate whether sertraline is a specific inhibitor of serine/glycine synthesis, we tested the efficacy of sertraline on cancer cell lines representing two subtypes of breast cancer, being those that rely on intracellular serine/glycine synthesis (breast cancer cell line MDA-MB-468) and those that depend on extracellular serine and glycine uptake (breast cancer cell line MDA-MB-231) (6,26). MDA-MB-468 cells typically express higher transcript and protein levels of serine/glycine synthesis enzymes (6,26), and flux through this metabolic pathway is associated with proliferation and sustained survival (6). We performed mass spectrometry based ^13^C_6_-glucose tracing experiments to monitor the activity of serine/glycine synthesis in more detail in these cell lines. In such experiments, synthesis of serine and glycine can be monitored by following carbon-13 stable isotope incorporation from glucose, with glucose-derived serine and glycine showing mass shifts of 3 and 2 units (M+3 serine and M+2 glycine), respectively. Cellular uptake of unlabeled serine and glycine from the cell culture medium will, on the other hand, result in unlabeled (M+0) serine and glycine, whereas interconversion between serine and glycine, catalyzed by SHMT1/2, will result in partially labeled serine (M+1 and M+2) and glycine (M+1) (Figure 3D). Our data confirmed that MDA-MB-468 cells showed a serine M+3 and glycine M+2 contribution from labeled glucose around 10%, while serine and glycine were unlabeled in MDA-MB-231 cells (Figure S2A). MDA-MB-468 cells are thus able to endogenously produce serine and glycine, whereas MDA-MB-231 cells are not. In agreement with this, real-time monitoring of cell confluency of both of these cell lines treated with NCT-503, an established PHDGH inhibitor (7), showed dose-dependent inhibition of MDA-MB-468 proliferation, while minimal effects on MDA-MB-231 were observed (Figure S2B-C). In accordance with these data, treatment of MDA-MB-468 cells with sertraline induced a dose-dependent impairment of the proliferation of these cells, whereas MDA-MB-231 cells were hardly affected (Figure 2 and S3). These results indicate that compounds re-sensitizing the lower eukaryotic model system, can specifically inhibit the proliferation of serine/glycine synthesis addicted breast cancer cells.

**Figure 2.**
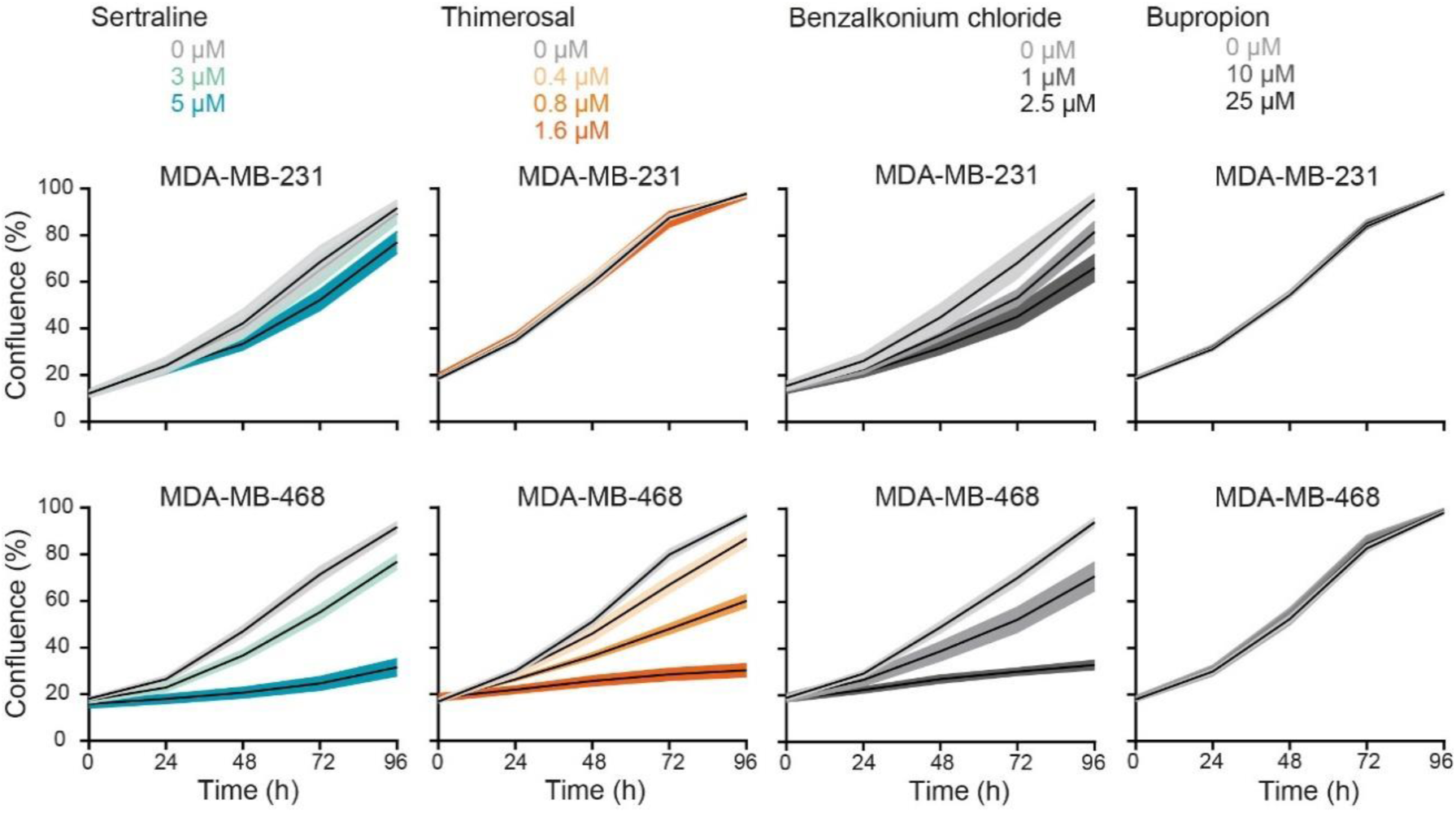
A subset of “re-sensitizing agents” impairs proliferation of serine/glycine synthesis addicted breast cancer cell lines. Proliferation during 96 h, as determined by real-time monitoring of cell confluence (%), of MDA-MB-231 (upper) and MDA-MB-468 (lower) cells upon treatment with indicated concentrations of sertraline, thimerosal, benzalkonium chloride and bupropion (left to right). 1 representative result of three biological replicates, containing each at least three technical replicates, is shown (mean ± SD).

**Figure 3.**
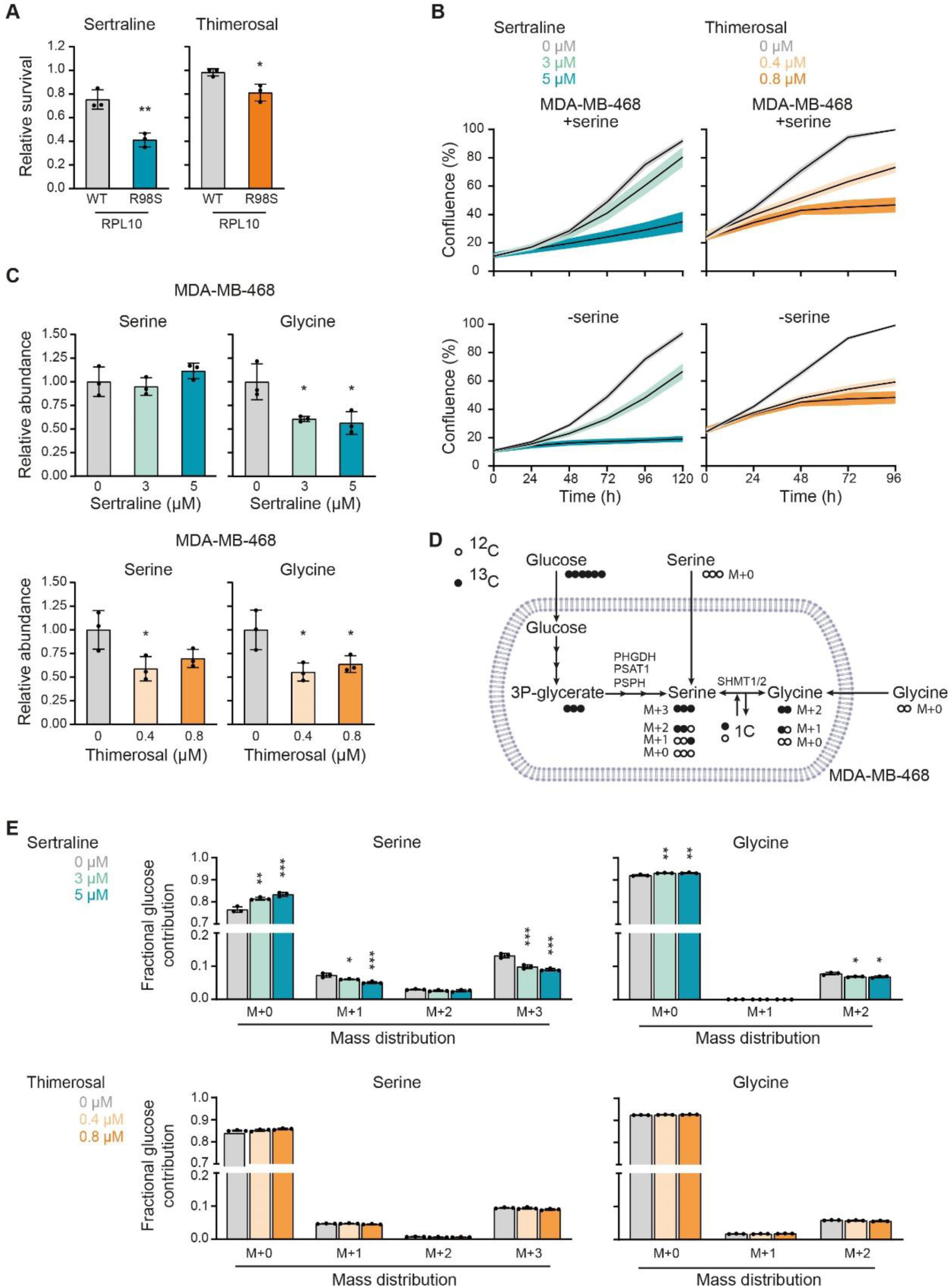
Sertraline and thimerosal target serine/glycine synthesis in cancer cells. **(A)** Survival, by measuring cell viability using flow cytometry, of Ba/F3 cells, that express either RPL10 WT or RPL10 R98S, upon treatment with 7.3 µM sertraline (left) or 1 µM thimerosal (right) for 48 h. Values are presented relative to the control treatment (n = 3 individual CRISPR/Cas9 clones with at least two technical replicates in each experiment, Student’s t-test). **(B)** Proliferation, as determined by real-time monitoring of cell confluence (%), of MDA-MB-468 cells cultured in DMEM with (upper) or without (lower) serine (400 µM) and treated with indicated concentrations of sertraline (left) or thimerosal (right). 1 representative result of three biological replicates, containing each at least three technical replicates, is shown. **(C)** Relative abundance of intracellular serine and glycine in MDA-MB-468 cells treated with indicated concentrations of sertraline (upper) and thimerosal (lower) for 72 and 24 h, respectively (n = 3, One-way ANOVA, Dunnett’s multiple comparisons test). **(D)** Schematic representation of carbon incorporation from ^13^C_6_-glucose into serine and glycine. Glucose-derived serine and glycine shows mass shifts of 3 and 2 units (M+3 serine and M+2 glycine), respectively. Cellular uptake of serine and glycine from the cell culture medium will result in unlabeled (M+0) serine and glycine, whereas interconversion between serine and glycine, catalyzed by SHMT1/2, will result in partially labeled serine (M+1 and M+2) and glycine (M+1). **(E)** Serine and glycine mass distribution showing the fractional glucose contribution of each mass upon treatment of MDA-MB-468 cells with indicated concentrations of sertraline (upper) or thimerosal (lower) (n = 3, One-way ANOVA, Dunnett’s multiple comparisons). In **(A-C and E)** data are presented as mean ± SD. *p < 0.05, **p < 0.01, ***p < 0.001.

Besides sertraline, a previously performed screening of 1600 off-patent drugs and other bioactive compounds (Pharmakon 1600 repurposing library) using the same yeast model system as described above, resulted in 49 other “re-sensitizing hits” (Table S1) (24,27). We hypothesized that this list might include other potent serine/glycine synthesis inhibitors (25). To select additional candidates for validation in serine/glycine synthesis addicted breast cancer cell lines, all 49 agents were ranked into three classes based on their yeast re-sensitizing capacity (high, intermediate and low). One representative agent of each class was retained for further validation, taking into account their clinical usage, as well as the presence of sulfhydryl containing groups that are proven to be effective for inhibition of PHGDH enzyme activity (8). Quaternary ammonium compounds came out as the top “hits” in the highest ranked class, of which we selected benzalkonium chloride. Out of the intermediate class of re-sensitizing agents, we selected thimerosal due to the presence of a reactive sulfhydryl group in its chemical structure. Bupropion, representing the lowest ranked class, was also retained for testing since it is clinically used as antidepressant, similar to sertraline. Thimerosal and benzalkonium chloride showed specific anticancer activity against the serine/glycine synthesis addicted MDA-MB-468 cells (Figure 2 and S3). Conversely, bupropion did not show any anticancer activity, not even at high concentrations (Figure 2 and S3).

To validate these results, we selected a second pair of breast cancer cell lines, of which one is dependent on serine/glycine synthesis (HCC70) and one favors the uptake of serine and glycine from the microenvironment (MCF7) (6,26). First, we tested the sensitivity to PHGDH inhibitor NCT-503. As expected, NCT-503 induced a stronger inhibition of proliferation in HCC70 than in MCF7, especially at lower drug doses (Figure S4). In agreement with previous results, sertraline, thimerosal and benzalkonium chloride impaired HCC70 cell proliferation in a dose-dependent manner, while there was no or minor toxicity on MCF7 cells (Figure S5). Overall, thimerosal showed the highest selectivity in targeting serine/glycine synthesis addicted breast cancer cells as compared to sertraline and benzalkonium chloride, while sertraline, as an antidepressant, has highest potential for clinical use in cancer patients. We therefore selected both compounds for further characterization of their mode-of-action.

### Sertraline and thimerosal target serine/glycine synthesis in cancer cells

Next, we verified whether sertraline and thimerosal would be applicable as agents that target serine/glycine synthesis addicted cancers other than breast cancer. The ribosomal RPL10 R98S mutation is detected in 8% of pediatric T-cell acute leukemia patients (28). In leukemic cells carrying the RPL10 R98S mutation, the mutant ribosomes accumulate on PSPH mRNAs, leading to elevated PSPH protein translation associated with enhanced serine/glycine synthesis. We showed that RPL10 R98S leukemic cancer cells depend on this elevated PSPH expression to support their proliferation and *in vivo* expansion (14). As such, RPL10 R98S leukemias represent a second cancer subgroup that is addicted to intracellular serine/glycine synthesis and we hypothesized that RPL10 R98S mutant cells might be more sensitive to sertraline and thimerosal. We used mouse pro-B Ba/F3 cells in which the endogenous Rpl10 locus was engineered with CRISPR/Cas9 technology to express the WT or R98S mutant form of RPL10, comparing three independent single cell clones of each genotype. RPL10 R98S clones showed selective inhibition of cell survival upon sertraline or thimerosal treatment as compared to the RPL10 WT clones (Figure 3A). These results confirm applicability of sertraline and thimerosal in serine/glycine synthesis addicted cancer types other than breast cancer and suggest that these agents may target different serine/glycine synthesis addicted cancer subtypes.

To look closer into the mode-of-action of both compounds and their potential impact on serine/glycine synthesis, we first evaluated whether serine depletion from the cell culture medium increases the sensitivity of MDA-MB-468 breast cancer cells to sertraline and thimerosal by enforced dependency on intracellular serine/glycine synthesis. Only sertraline induced a significantly stronger impairment of MDA-MB-468 cell proliferation under serine-depleted conditions (Figure 3B and S6A-B). A similar trend was observed with NCT-503 (Figure S6C-D).

Given that MDA-MB-468 cells rely more on synthesis than uptake, inhibition of serine/glycine synthesis could cause changes in intracellular serine and/or glycine abundance, depending on the specific synthesis enzyme that is targeted. While reductions in enzymatic activity of PHGDH, PSAT1 or PSPH are expected to result in reduced intracellular concentrations of both serine and glycine, targeting SHMT1/2 is rather expected to affect levels of glycine only. Mass spectrometry analysis revealed that sertraline did not affect the intracellular levels of serine in MDA-MB-468 cells, and that it solely reduced the abundance of downstream glycine (Figure 3C). In contrast, thimerosal decreased serine abundance to the same extent as glycine, as seen for NCT-503 (Figure 3C and S6E).

We subsequently performed mass spectrometry based ^13^C_6_-glucose tracing to monitor the effects of sertraline and thimerosal on overall serine/glycine synthesis in more detail (Figure 3D). Our data showed that sertraline reduced *de novo* synthesized M+3 serine, and that it targets serine to glycine interconversion in MDA-MB-468 cells, characterized by a reduced fractional labeling of M+2 glycine and M+1 serine (Figure 3E). In contrast to sertraline, thimerosal did not significantly reduce the fraction of glucose-derived M+3 serine and M+2 glycine (Figure 3E). In parallel, NCT-503 only caused minor changes in fractional glucose contribution of M+3 serine and M+2 glycine (Figure S6F) that could not solely account for the lower intracellular serine and glycine abundances (Figure S6E). Since we assess with this tracing approach the ratio between synthesis and uptake, our data may indicate a simultaneous downregulation of serine/glycine synthesis and uptake upon thimerosal or NCT-503 treatment.

Putting these results together, we reasoned that thimerosal, similar to NCT-503, is more likely to affect one of the three first biosynthetic enzymes. For sertraline, our data point towards inhibition of serine to glycine interconversion, implying that SHMT1/2 can be its potential target. In line with this hypothesis, steady state levels of serine (Figure 3C) may originate from a block in conversion to glycine and subsequent downregulation of *de novo* serine synthesis and uptake.

### Thimerosal reduces PHGDH activity, while sertraline directly inhibits downstream SHMT

To gain additional insights in the exact serine/glycine synthesis enzyme targeted by sertraline and thimerosal, we first performed an *in vitro* enzymatic PHGDH assay. Sertraline did not show any effect on *in vitro* PHGDH enzymatic activity, further excluding PHGDH as sertraline’s potential target (Figure 4A). Similar to NCT-503, thimerosal was able to reduce *in vitro* PHGDH enzymatic activity (Figure 4A). Notably, the thimerosal concentration inducing 50% inhibition (IC50 = 0.1517 µM) was even 10-fold lower than this of NCT-503 (IC50 = 1.747 µM) (Figure 4A and S7A).

**Figure 4.**
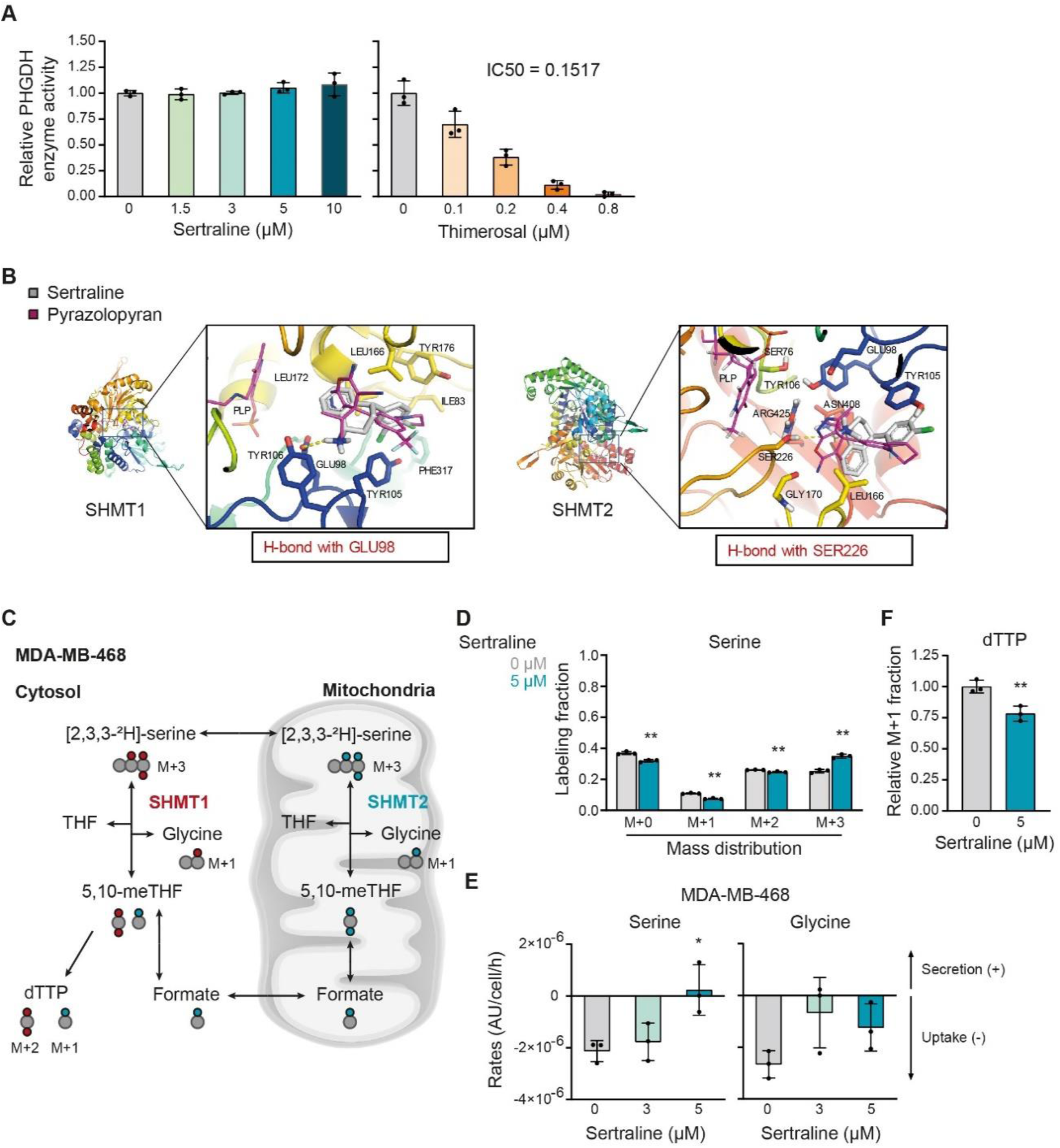
Thimerosal reduces PHGDH activity, while sertraline directly targets downstream SHMT. (A) PHGDH (= phosphoglycerate dehydrogenase) enzymatic *in vitro* assay, measuring PHGDH activity upon addition of indicated concentrations of sertraline (left) or thimerosal (right). Values are presented relative to the control (n = 3). **(B)** SHMT1 (left) and SHMT2 (right) in complex with sertraline (grey), with a magnified view of the binding pocket showing the interactions formed by sertraline. H-bonds formed by sertraline are presented as yellow dashes. The known SHMT inhibitor, with a pyrazolopyran scaffold, is shown in magenta. **(C)** Schematic overview of isotopic tracing with [2,3,3-^2^H]-serine, showing ^2^H incorporation in downstream metabolites glycine and thymidine (dTTP). Cells taking up fully deuterated (M+3) serine use this to synthesize glycine with one deuterium label (M+1). While cytosolic methylene-THF production, by SHMT1, results in double ^2^H-labeled (M+2) dTTP (red dots), mitochondrial SHMT2 will generate single ^2^H-labeled (M+1) dTTP (blue dots). **(D)** Serine mass distribution showing the labeling fraction of each mass upon treatment of MDA-MB-468 cells with control (DMSO) or sertraline (5 µM) for 48 h (n = 3, Multiple t-test). **(E)** Serine and glycine uptake (-) and secretion (+) rates (AU/cell/h) of MDA-MB-468 cells treated with indicated concentrations of sertraline for 72 h (n = 3, One-way ANOVA, Dunnett’s multiple comparisons). **(F)** Deuterium M+1 labeled dTTP fraction in MDA-MB-468 cells treated with control (DMSO) or sertraline (5 µM) for 48 h. Values are presented relative to the control (n = 3, Student’s t-test). In **(A and D-F)** data are presented as mean ± SD. *p < 0.05, **p < 0.01.

As our *in vitro* enzymatic PHGDH assay had excluded PHGDH as being sertraline’s target (Figure 4A) and previous data already pointed towards more downstream inhibition of serine/glycine synthesis (Figure 3), we focused further on SHMT1/2. To do this, we computationally docked sertraline to both SHMT1 and SHMT2. It has been shown that SHMT is a ubiquitous pyrodoxal 5’-phosphate(PLP)-dependent enzyme (17,29). Therefore, docking scores were determined with the PLP co-factor inside the binding pocket (Table 1). As a reference, the already reported plant derived and non-clinically used dual SHMT1/2 inhibitor with a pyrazolopyran scaffold, was used (17,30). High docking scores were obtained for sertraline, in the range of the compounds characterized by a pyrazolopyran scaffold. Moreover, aligning both dockings showed that sertraline potentially binds in the exact same pocket as the pyrazolopyran scaffold and has similar interactions with SHMT1/2, namely a hydrogen bond with its -NH group (Figure 4B).

**Table 1.** Computational docking of sertraline to SHMT1 and SHMT2. Docking scores for sertraline into the active pocket of SHMT1 and SHMT2. The already identified dual SHMT inhibitor, based on a pyrazolopyran scaffold, was used as reference structure.

| Interaction | Docking score |
| --- | --- |
| SHMT1_sertraline1<sup>a</sup> | 61.49 |
| SHMT1_sertraline2<sup>a</sup> | 59.68 |
| SHMT1_pyrazolopyran (=reference) | 66.65 |
| SHMT2_sertraline | 51.34 |
| SHMT2_pyrazolopyran (=reference) | 65.39 |

<sup>a</sup> 1 and 2 refer to the two conformations in which sertraline can bind the SHMT1.

To experimentally measure cellular SHMT activity upon sertraline treatment, we performed an isotopic hydrogen tracer analysis (Figure 4C) in which MDA-MB-468 breast cancer cells were incubated for 48 hours with [2,3,3-^2^H]-serine, followed by analysis of deuterated 2H incorporation in downstream metabolites glycine and thymidine triphosphate (dTTP) (31). These tracing studies revealed that sertraline increased the intracellular fraction of M+3 serine and decreased the fraction of M+0, M+1 and M+2 serine (Figure 4D). While M+3 serine is used by SHMT1/2 to convert to glycine, partially labelled serine (M+1 and M+2) only arises from converting glycine back into serine. As such, these findings together imply a reduced conversion flux between serine and glycine and might further support a block in general SHMT activity.

In contrast to what may be expected from an SHMT inhibitor, sertraline did not increase the intracellular serine levels (Figure 3C). To look closer into this, we determined the impact of sertraline on serine and glycine uptake and secretion rates in MDA-MB-468 cells (Figure 4E). Our data showed a dose-dependent reduction of net serine uptake. Upon the highest dose of sertraline, we even observed net serine secretion. These observations can explain the lack of serine accumulation in these breast cancer cells. Whether these effects are due to direct secretion of serine and/or a sensory feedback mechanism of pyruvate kinase (PKM2), an important regulator of the glycolytic flux, remains to be determined. Furthermore, measuring glycine uptake and secretion rates revealed that sertraline-treated MDA-MB-468 cells have the tendency to take up less glycine (Figure 4E). The strength of blocking both the interconversion of serine into glycine and glycine uptake is what makes SHMT inhibitors more cytotoxic, explaining sertraline’s potent activity on MDA-MB-468 cells, as was previously observed using the known, but not clinically used, dual SHMT1/2 inhibitor (17). Collectively, these results support that sertraline inhibits SHMT activity and glycine uptake, leading to reduced net serine uptake and intracellular *de novo* serine synthesis, to prevent serine accumulation.

To distinguish between cytosolic SHMT1 and mitochondrial SHMT2 activity, we evaluated deuterated serine incorporation into dTTP (Figure 4C). While cytosolic SHMT1 will produce M+2 dTTP, mitochondrial SHMT2 will generate M+1 dTTP via transfer of one-carbon (1C) units from the mitochondria to the cytosol (31,32). In proliferating MDA-MB-468 breast cancer cells, only M+1 dTTP was detected (Figure S7B), supporting that these cells process their 1C units only in the mitochondria via SHMT2. Interestingly, the dTTP fraction with one deuterium label (M+1) was decreased upon sertraline treatment (Figure 4F). Therefore, we concluded that sertraline inhibits thymidine synthesis by blocking SHMT2 activity, thereby targeting cancer cell proliferation.

### Sertraline has clinical potential, especially when used in combination therapy

In contrast to thimerosal, for which clinical applications are limited to topical use because of a toxic mercury group in its structure, sertraline is a small molecule serotonin blocker that is widely used in the clinic as antidepressant. Our findings support that sertraline can be an attractive adjuvant therapeutic agent to specifically treat serine/glycine synthesis addicted cancers.

It has been established that mitochondrial dysfunction causes changes in one-carbon metabolism and that cancer cells will depend even more on their serine/glycine synthesis upon mitochondrial inhibition (33,34). Furthermore, serine/glycine synthesis and the tricarboxylic acid (TCA) cycle are strongly interconnected, implying that targeting both pathways might further metabolically disrupt cancer cells. Since sertraline targets serine/glycine synthesis, combining it with drugs causing mitochondrial dysfunction might strengthen its anticancer activity. As a proof-of-concept, we investigated the effect of sertraline on MDA-MB-468 proliferation when combined with respiratory chain inhibitors rotenone and antimycin A, targeting complex I and complex III, respectively (35). To test this, we used sub-optimal dosages of rotenone and antimycin A. While single drug treatment with sertraline caused around 20% reduction in cell proliferation, the combination of sertraline with both rotenone and antimycin A indeed further decreased proliferation of MDA-MB-468. No synergistic effect of both combinations on MDA-MB-231 proliferation was observed (Figure 5A and S8A-C).

**Figure 5.**
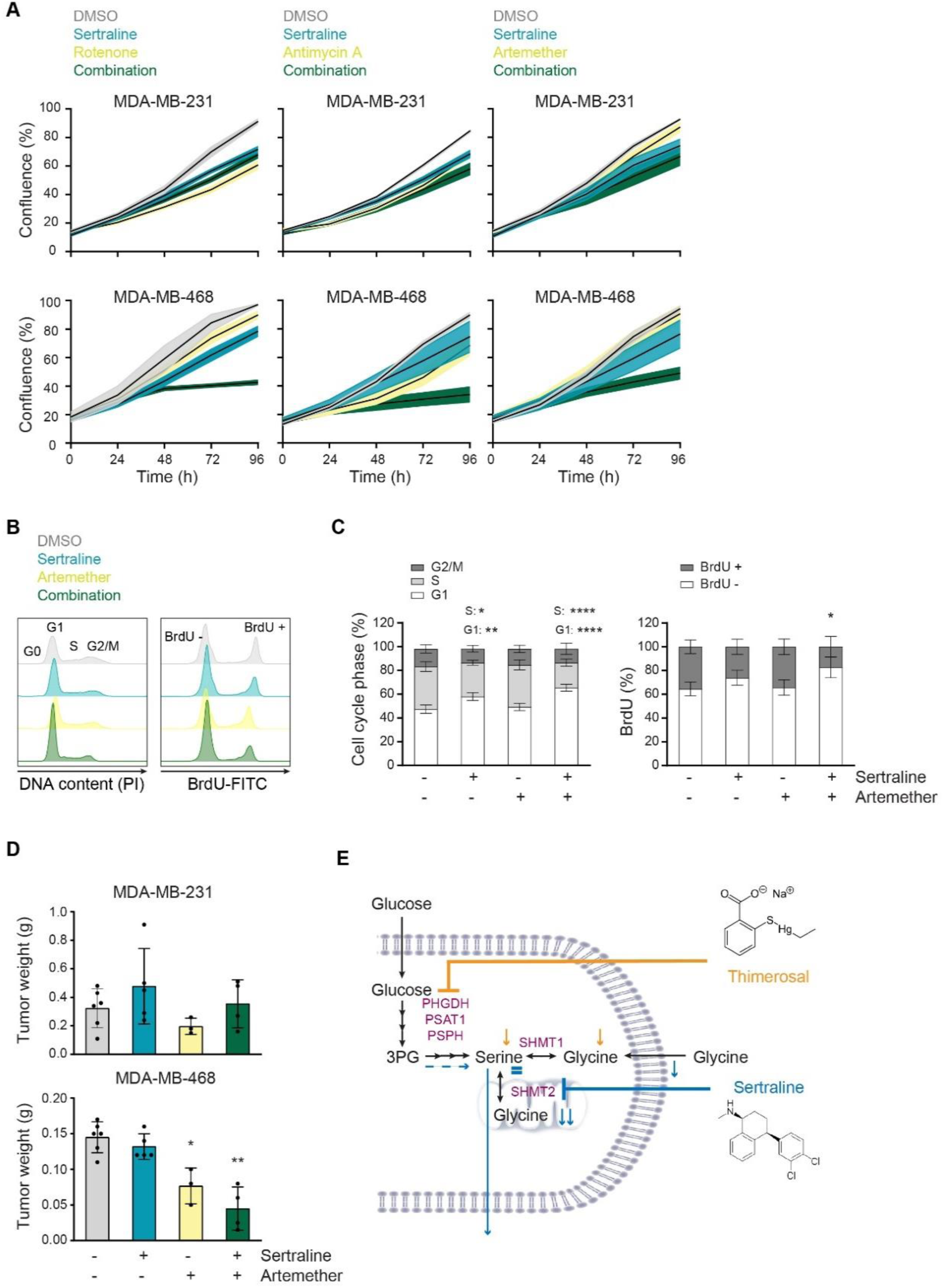
Sertraline has clinical potential, especially in combination with mitochondrial inhibitors. **(A)** Proliferation during 96 h, as determined by real-time monitoring of cell confluence (%), of MDA-MB-231 (upper) and MDA-MB-468 (lower) cells upon treatment with sertraline (5 µM) in combination with rotenone (50 nM), antimycin A (50 nM) or artemether (80 µM). 1 representative result of three biological replicates, containing each at least three technical replicates, is shown. **(B)** Histograms showing PI cell cycle analysis (left) and BrdU incorporation (right) of MDA-MB-468 cells treated with DMSO, sertraline (5 µM) and/or artemether (80 µM) for 24 h. 1 representative result of three biological replicates is shown. **(C)** Quantification of (B) pooling all three biological replicates (n = 3, Two-way ANOVA, Dunnett’s multiple comparisons test). **(D)** Tumor weight (g) of MDA-MB-231 (left flank) and MDA-MB-468 (right flank) mouse xenografts after treatment with DMSO, sertraline (2,5 mg/kg), artemether (40 mg/kg) or a combination of both compounds for 4 weeks (n ≥ 3, Kruskal-Wallis test, followed by Mann-Whitney U test). **(E)** Schematic overview of the mode-of-action of sertraline and thimerosal. In **(A and D)** data are presented as mean ± SD. *p< 0.05, **p < 0.01.

To further analyze the clinical potential of sertraline in combination therapy, we tested sertraline in combination with a clinically used and less generally toxic drug than rotenone and antimycin A. Artemether, an antimalarial agent, has already been shown to have potent anticancer activity (36,37). Interestingly, artemisinin, which belongs to the same structural class of natural plant compounds as artemether, was identified as activator of AMPK, suggesting a possible inhibition of mitochondrial function (38). By performing metabolic ^13^C_6_-glucose tracing of artemether-treated MDA-MB-468 cells (Figure S9), we confirmed inhibitory action of artemether on mitochondrial TCA cycle activity, as evidenced by decreased contribution of labeled glucose into TCA cycle metabolites, especially the M+4 variants. Besides, a slight reduction of the M+3 serine fraction was detected, but this was lower than the observed inhibitory effect on TCA cycle intermediates. Additionally, glucose contribution into M+3 lactate and M+3 alanine was elevated upon artemether treatment, which was not observed with sertraline. The latter suggests that overall TCA cycle activity is reduced, and pyruvate will use alternative routes such as its reduction to lactate, supporting that artemether acts as mild mitochondrial inhibitor. Corresponding to what was observed using rotenone and antimycin A, the combination of suboptimal dosages of sertraline and artemether indeed further decreased proliferation of MDA-MB-468 as compared to single drug treatment, while minimal effects where observed on MDA-MB-231 (Figure 5A and S8B-C). Metabolic ^13^C_6_-glucose tracing confirmed the superior effect of the novel combination therapy (Figure S9), as the contribution of labeled glucose into TCA cycle metabolites was stronger reduced. In contrast, the reduction in M+3 serine and M+2 glycine was identical to this of sertraline treatment alone, confirming that the inhibitory effect on serine/glycine synthesis is rather due to sertraline than to artemether.

Since the artemether-sertraline combination mainly reduced the MDA-MB-468 proliferation rate, we hypothesized that it interferes with cell cycle progression rather than inducing cell death. Indeed, bromodeoxyuridine (BrdU) staining showed a significant decrease in BrdU incorporation in MDA-MB-468 cells after 24 hours of combination treatment, compared to control and single drug treatment (Figure 5B-C and S10B). Moreover, propidium iodide (PI) cell cycle analysis confirmed a G1-S cell cycle arrest, with 20% more MDA-MB-468 cells in G1 phase upon combination treatment (Figure 5B-C and S10B). Only 5% of MDA-MB-468 cells were double positive for the Annexin V and Zombie Aqua cell death markers after 24 hours treatment with the combination (Figure S10C), supporting limited effects on cell apoptosis. Apart from the artemether-sertraline combination, minor effects on cell cycle were observed with a sub-optimal dose of sertraline that increases with increased doses of sertraline (Figure S10A-B). As such, artemether seems to strengthen sertraline’s inhibitory effect on G1-S cell cycle progression, highlighting the potential of a mitochondrial inhibitor in combination with sertraline.

Finally, to evaluate the therapeutic potential of these findings more profoundly, we tested the efficacy of the artemether-sertraline combination in an *in vivo* mouse model. To this end, we implanted MDA-MB-231 and MDA-MB-468 cell lines in the opposite flanks of immunodeficient NOD-SCID/IL2γ-/- (NSG) mice and treated the animals with DMSO, sertraline (2.5 mg/kg), artemether (40 mg/kg) or a combination of the two compounds. After 4 weeks, only the MDA-MB-468 mouse xenografted tumors showed significant differences between treatment groups and artemether significantly reduced MDA-MB-468 tumor growth (Figure 5D). Most notably, the combination of sertraline and artemether caused an even stronger inhibition of MDA-MB-468 tumor formation *in vivo* as compared to monotherapy (Figure 5D). No changes in *in vivo* MDA-MB-231 tumor growth were observed between treatment groups within the same mice (Figure 5D). In conclusion, the combination of sertraline and artemether not only reduces *in vitro* proliferation of serine/glycine synthesis addicted breast cancer cell lines, but is also effective in limiting *in vivo* breast cancer xenograft growth depending on intracellular serine/glycine synthesis. Collectively, combining a serine/glycine synthesis inhibitor, such as sertraline, with drugs targeting mitochondrial function seems a valuable treatment strategy for serine/glycine synthesis addicted cancers.

## DISCUSSION

Our data support upregulation of serine/glycine synthesis as a general mechanism to deal with stress and/or to support cell growth that has been evolutionary conserved from the lower eukaryote yeast *C. albicans* up to human cancer cells. Hence, this yeast model system can be used for rapid and low-cost screening for compounds targeting serine/glycine synthesis. Further characterization of the mode-of-action of “re-sensitizing hits” revealed that sertraline and thimerosal target intracellular serine/glycine synthesis by reducing the enzymatic activity of SHMT2 and PHGDH, respectively (Figure 5E). Yet, we cannot exclude sertraline’s inhibitory activity towards SHMT1. Interestingly, it has recently been established that both SHMT1 and SHMT2 need to be suppressed to inhibit proliferation of serine/glycine synthesis addicted T-ALL cell lines (23). Furthermore, molecular docking supported binding of sertraline in the active pocket of SHMT1 (Figure 3B), that was comparable with binding of the already identified non-clinical dual SHMT1/2 inhibitor (17,30).

Although thimerosal showed potent PHGDH inhibition with 10-fold lower IC50 values compared to NCT-503, its clinical applications are limited to topical use due to the presence of a toxic mercury group in its chemical structure (Figure 5E). Therefore, we tested thimerosal derivatives that do not contain a mercury group, such as methyl thiosalicylate, thiosalicylic acid and sodium thiosalicylate, for their potential to specifically inhibit MDA-MB-468 proliferation. However, these compounds did not affect the proliferation of MDA-MB-468 cells, nor could they inhibit *in vitro* PHGDH enzymatic activity. In agreement with these observations, it was previously described that thimerosal’s sulfhydryl reactive properties, which are expected to be important for inhibiting PHGDH activity, are attributable to its mercury-containing chemical structure (8,39). In line with an important role for sulfhydryl reactive properties in inhibiting PHGDH activity, compound CBR-9480, being very similar in chemical structure with thimerosal, was also picked up as a potent inhibitor of PHGDH (8). In contrast to thimerosal, the antidepressant sertraline has high clinical potential as targeting agent for serine/glycine synthesis addicted cancers.

Besides T-ALL (14,23) and breast cancer, upregulation of serine/glycine synthesis is also known in other types of cancer such as melanoma, glioblastoma, prostate, testis, ovary, liver, kidney, lung and pancreas cancer (40,41). Expression levels of serine/glycine synthesis enzymes reaching above 4-fold increased expression as compared to normal tissue controls (based on our Ba/F3 RPL10 WT and R98S model), may serve as a biomarker to identify serine/glycine synthesis addicted cancers that may benefit from sertraline treatment.

Whereas antitumor activity of sertraline has previously been described (42–44), our findings pinpoint the exact mode-of-action of its anticancer activity and support that this agent could also be an attractive adjuvant therapeutic agent to specifically treat the subset of serine/glycine synthesis addicted cancers. Moreover, suppression of serine/glycine synthesis has been shown to re-sensitize therapy resistant tumors to anticancer treatment. In particular, PHGDH targeting resulted in re-sensitization to doxorubicin, vemurafenib and HIF2α-antagonist in therapy resistant triple-negative breast cancer cells, melanoma with oncogenic BRAF V600E mutations and advanced renal cell carcinoma, respectively (45–47). In addition, sertraline passes the blood-brain barrier, which may even open opportunities to target serine/glycine synthesis addicted brain tumors.

The doses at which we observed activity against serine/glycine synthesis addicted cancer cell lines are in the range of concentrations at which sertraline’s antitumor activity was described by others (42–44). Clinical use of sertraline as anticancer adjuvant therapeutic compound will only be possible if the dosages required for anticancer efficacy are not reaching toxicity in humans. As an antidepressant, sertraline is administered in a dosage ranging from 50 mg to 200 mg a day, resulting in sertraline serum concentrations between 0.065 and 0.54 µM (48). Considering that the average body weight of an adult in Europe is 70.8 kg (49), sertraline doses vary between 0.7 and 2.8 mg/kg. Furthermore, sertraline has a linear pharmacokinetic profile and daily oral intake of a dose of 400 mg (5.6 mg/kg), resulting in a plasma concentration of 0.82 µM, is well tolerated in patients (43,48). In our *in vivo* experiments, we treated the mice with 2.5 mg/kg sertraline, which thus is in the range of the administered doses in humans. Hassell *et al*. (2016) even used doses of 60 mg/kg in mice and no adverse effects were observed (43). This suggest that therapeutics doses of sertraline can be achieved in cancer patients.

In conclusion, we identified the widely used antidepressant sertraline as a novel inhibitor of serine/glycine synthesis enzyme SHMT that demonstrated high selectivity and efficacy against metabolically addicted cancers. Whereas previously identified inhibitors of serine/glycine synthesis enzymes did not reach clinical trials, sertraline is a clinically used drug that can safely be used in humans. Our results suggest that sertraline may have applications as adjuvant therapeutic intervention strategy for serine/glycine synthesis addicted cancers, especially when combined with drugs attacking other central nodes of cancer cell metabolism such as mitochondrial inhibitors. Collectively, this work provides a novel and cost-efficient treatment option for the rapidly growing list of serine/glycine synthesis addicted cancers.

## ACKNOWLEDGEMENTS

The authors would like to thank Annelies Peeters and Tamara Davenne for technical support.

