## Supplemental file for "Repurposing the antidepressant sertraline as SHMT inhibitor to suppress serine/glycine synthesis addicted breast tumor growth"

### ADDITIONAL FILE FOR:

[1]Laboratory for Disease Mechanisms in Cancer, Department of Oncology, KU Leuven and Leuven Cancer Institute (LKI), Herestraat 49, 3000 Leuven, Belgium; [2]Centre of Microbial and Plant Genetics – Plant Fungi Interactions (CMPG-PFI), KU Leuven, Kasteelpark Arenberg 20, 3001 Heverlee, Belgium; [3]Maastricht University Medical Center, Department of Radiation Oncology (MAASTRO), GROW School for Oncology and Developmental Biology, Maastricht, The Netherlands; [4]Laboratory of Cellular Metabolism and Metabolic Regulation, VIB-KU Leuven Center for Cancer Biology, VIB Leuven, Herestraat 49, 3000 Leuven, Belgium; [5]Laboratory of Cellular Metabolism and Metabolic Regulation, Department of Oncology, KU Leuven and Leuven Cancer Institute (LKI), Herestraat 49, 3000 Leuven, Belgium; [6]Department of Chemistry, KU Leuven, Celestijnenlaan 200G, 3001 Heverlee, Belgium; [7]Department of Cardiovascular Sciences, University Hospitals Leuven, Herestraat 49, 3000 Leuven, Belgium; [8]Department of Hepatology, University Hospitals Leuven, Herestraat 49, 3000 Leuven, Belgium; [9]Department of Plant Systems Biology, VIB, Technologiepark 927, 9052 Gent, Belgium.

[†]Shared first authors

[‡]Shared last authors

**Running title**: Sertraline targets serine/glycine synthesis enzyme SHMT.

**Keywords:** Breast cancer, Cancer metabolism, Serine/glycine synthesis, Sertraline, SHMT

**Financial support:**

SLG received a SB PhD fellowship from “Fonds Wetenschappelijk Onderzoek (FWO) (1S14517N). KRK was supported by a PDM postdoctoral mandate fellowship obtained from the KU Leuven, the

“Emmanuel van der Schueren” postdoctoral fellowship from “Kom op tegen Kanker” and a research grant from FWO (FWO KAN2018 1501419N). GR is supported by consecutive PhD fellowships from the “Emmanuel van der Schueren - Kom op tegen Kanker” foundation and FWO (1137117N and 1137119N). PG and AV acknowledge a research grant from FWO (G0F9316N Odysseus). SV and BV are SB PhD fellow at FWO (1S07118N and 1S49817N). PV and DC are senior clinical investigators of the Research Foundation - Flanders. SMF acknowledges FWO funding and KU Leuven Methusalem co-funding. KT received a mandate as Innovation Manager from IOF Fund of KU Leuven. This research was funded by a grant from “Stichting Tegen Kanker” (2016-112) and by funding from the KU Leuven Research Council (C1 grant C14/18/104) to KDK.

**Corresponding authors:**

* Kim De Keersmaecker, KU Leuven, Laboratory for Disease Mechanisms in Cancer, Department of Oncology, KU Leuven and Leuven Cancer Institute (LKI), Campus Gasthuisberg O&N1, box 603, Herestraat 49, 3000 Leuven. Phone number: +32 16 37 31 67.


* Karin Thevissen, KU Leuven, Centre of Microbial and Plant Genetics – Plant Fungi Interactions (CMPG-PFI), Kasteelpark Arenberg 20, box 2460, 3001 Heverlee. Phone number: +32 16 32 96 88 or +32 16 32 16 31.


**Conflict of interest:**

SMF has received funding from Bayer, Merck and Black Belt Therapeutics. All other authors declare no potential conflicts of interest.

**This PDF file includes:** Supplementary Note
Supplementary Methods
Table S1
Figures S1-S10
SI References

#### SUPPLEMENTARY NOTE

##### A serine/glycine synthesis dependent lower eukaryotic yeast model

The yeast *Candida albicans*, the major fungal pathogen for humans, typically forms drug-tolerant biofilms on biotic surfaces such as the skin, the mouth, the human gastro-intestinal tract and genital area. To treat this type of mucosal fungal infection, azoles, such as miconazole, are currently used. In contrast to free-living *C. albicans* cells, the increasing tolerance of biofilm cells to azoles makes biofilm-associated infections hard to eradicate (1). Specifically, biofilm cells are up to thousand-fold more tolerant to miconazole than their planktonic counterparts (2).

Revisiting previously obtained transcriptome data of miconazole-induced tolerance pathways in *C. albicans* biofilm cells highlighted a biofilm-specific upregulation of genes involved in serine/glycine synthesis (*SHM2*) and one-carbon metabolism (*MET13, SAH1, GCV1, GCV2* and *MIS11*) (3). In support of these data, conditioned medium of *C. albicans* biofilms that were treated with a sublethal dose of miconazole contained an excess of serine and glycine as compared to conditioned medium of control cultures (Figure S1A). In this experimental set-up, there was no difference in reductive potential between miconazole-stressed *C. albicans* biofilm cells and control cells (Figure S1B). We therefore reasoned that miconazole-stressed *C. albicans* biofilm cells upregulate their serine/glycine synthesis as a tolerance mechanism against sublethal miconazole doses. Hence, miconazole-stressed *C. albicans* biofilms can serve as a valuable platform to screen for compounds targeting serine/glycine synthesis, which are expected to re-sensitize the biofilms to miconazole.

#### SUPPLEMENTARY MATERIAL AND METHODS

***C. albicans* yeast strain and chemicals:** *C. albicans* strain SC5314 (4) was grown routinely on YPD (1% yeast extract, 2% peptone (International Medical Products) and 2% glucose (Sigma-Aldrich)) agar plates at 30 °C. RPMI-1640 medium (pH 7.0) (Sigma-Aldrich) with L-glutamine and without sodium bicarbonate was buffered with MOPS (Sigma-Aldrich). Stock solutions of miconazole (Sigma-Aldrich) and sertraline (Sigma-Aldrich) were prepared in dimethyl sulfoxide (DMSO) (VWR International) and stored at -20 °C.

***C. albicans* serine and glycine medium measurements:** A *C. albicans* SC5314 overnight culture, grown in YPD, was diluted to an optical density (OD) of 0.1 (approximately $10^6$ cells/ml) in RPMI-1640 medium (Sigma-Aldrich) and 2 ml of this suspension was added to the wells of a 6-well plate (Greiner Bio-One). After 1 h of adhesion at 37 °C, the medium was aspirated and biofilms were washed with phosphate buffered saline (PBS) to remove non-adherent cells, followed by addition of 2 ml fresh RPMI-1640 medium. Subsequently, the biofilms were grown for 24 h at 37 °C. After washing the biofilms with PBS, miconazole (75 µM) was added in RPMI-1640, resulting in a DMSO background of 0.2%. Next, biofilms were incubated for an additional 24 h at 37 °C. Finally, 1 ml of conditioned medium was collected and serine and glycine were quantified by cation-exchange chromatography on a Biochrom 30 analyzer (Biochrom, Cambourne, UK).

***C. albicans* metabolic activity assay:** Biofilms were grown in 6-well plates (Greiner Bio-One) and treated in RPMI-1640 medium (Sigma-Aldrich) as described above. After washing with PBS, 2 ml Cell-Titer Blue (CTB; Promega) (5), diluted 1/100 in PBS, was added to each well. After 1 h of incubation in the dark at 37 °C, fluorescence was measured with a fluorescence spectrometer (Synergy Mx Multimode Microplate Reader; BioTek) at $\lambda_{ex}$ of 535 nm and $\lambda_{em}$ of 590 nm. Finally, the percentage of metabolically active biofilm cells was calculated as described before (6).

***C. albicans* nuclear membrane permeability assay:** Biofilms were grown in 96-well plates and treated with miconazole (75 µM) and/or sertraline (75 µM) in RPMI-1640 medium. After washing with PBS, propidium iodide staining (Sigma-Aldrich) was performed as previously described (7).

**SUPPLEMENTARY TABLES**

**Table S1. List of yeast re-sensitizing agents.**

| Re-sensitizing capacity | Compound | Clinical use |
|---|---|---|
| **HIGH** (>90%) | Clioquinol | Antifungal, antiprotozoal |
| | Benzalkonium chloride | Disinfectant, antimicrobial, antifungal |
| | Dequalinium chloride | Antiseptic, disinfectant, bacteriostatic |
| | Methylbenzethonium chloride | Antiseptic, disinfectant, antimicrobial |
| | Pyrvinium pamoate | Anthelmintic |
| | Hexachlorophene | Disinfectant, antibacterial, antifungal |
| | Dichlorophene | Antifungal, antimicrobial, anticestodal |
| | Bithionate sodium | Anthelmintic |
| | Alexidine hydrochloride | Antimicrobial |
| | Gentian/Crystal violet | Antibacterial, antifungal, anthelmintic |
| **INTERMEDIATE** (50-90%) | Thimerosal **(sulfur groups)** | Antiseptic, antifungal |
| | Phenylmercuric acetate | Preservative, disinfectant |
| | Nitroxoline | Antibacterial |
| | Chloroxine | Antibacterial |
| | Piroctone olamine | Antifungal, anti-dandruff shampoo |
| | Ciclopirox olamine | Antifungal |
| | Pyrithione zinc **(sulfur groups)** | Fungistatic, bacteriostatic |
| | Broxyquinoline | Antiprotozoal |
| | Iodoquinol | Treatment of amoebiasis |
| | Evans blue | Viability assay dye |
| | Artemisinin | Antimalarial |
| | Artenimol | Antimalarial |
| | Abamectin | Insecticide, anthelmintic |
| **LOW** (<50%) | Suloctidil **(sulfur groups)** | Vasodilator |
| | Isosorbide mononitrate | Treatment of heart related chest pain |
| | Sodium nitroprusside | Antihypertensive |
| | Temozolomide | Oral chemotherapy drug |
| | Folic acid | Vitamin $B_9$ |
| | Bupropion | Antidepressant |
| | Sulfamethoxypyridazine | Antibacterial |
| | Oxantel pamoate | Anthelmintic |
| | Tioconazole **(sulfur groups)** | Antifungal |
| | Butoconazole **(sulfur groups)** | Antifungal |
| | Flucytosine | Antifungal |
| | Diacetamate | Analgesic, anti-inflammatory drug |
| | Artemether | Antimalarial |
| | Artesunate | Antimalarial |
| | Orbifloxacin | Antibacterial |
| | Salinomycin | Antibacterial |
| | Methenamine | Hardening component, binder |
| | Candicidin | Antifungal |
| | Warfarin | Anticoagulant, blood thinner |
| | Carvedilol phosphate | Antihypertensive |
| | Acipimox | Lipid-lowering agent |
| | Merbromin | Antiseptic |
| | Phentermine | Treatment of obesity |
| | Malathion **(sulfur groups)** | Insecticide |
| | Methylene blue **(sulfur groups)** | Treatment of methemoglobinemia, dye |
| | Sulfanilate zinc **(sulfur groups)** | Antibacterial |

**SUPPLEMENTARY FIGURES**

**Figure S1**

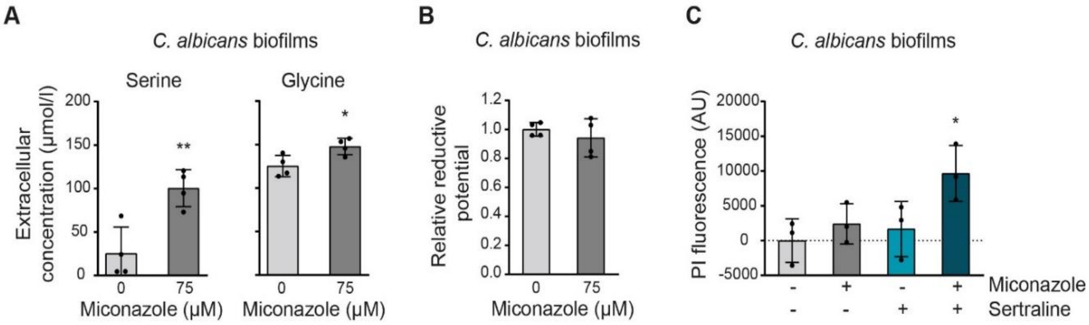


**Figure S1. Sertraline re-sensitizes *Candida albicans* biofilms that upregulate their serine/glycine synthesis as a tolerance mechanism against sublethal antifungal stress. (A)** Serine (left) and glycine (right) extracellular concentrations (µmol/l) of *C. albicans* biofilms treated with control (DMSO) or miconazole (75 µM) for 24 h (n = 4, Student's t-test). **(B)** Reductive potential, measured with Cell-Titer Blue, of miconazole-stressed (75 µM) *C. albicans* biofilm cells and control cells (DMSO) (n = 4, Student's t-test). **(C)** Cell death, measured using propidium iodide (PI), of *C. albicans* biofilms treated with control (DMSO), miconazole (75 µM), sertraline (75 µM) or a combination of both (n = 3, Two-way ANOVA, Dunnett's multiple comparisons test). While 75 µM of miconazole or sertraline monotherapy was not significantly affecting cell death, sertraline was able to re-sensitize miconazole's activity, increasing cell death of *C. albicans* biofilm cells by almost 4-fold compared to single drug treatment. In **(A-C)** data are presented as mean ± SD. *$p < 0.05$, **$p < 0.01$.

**Figure S2**

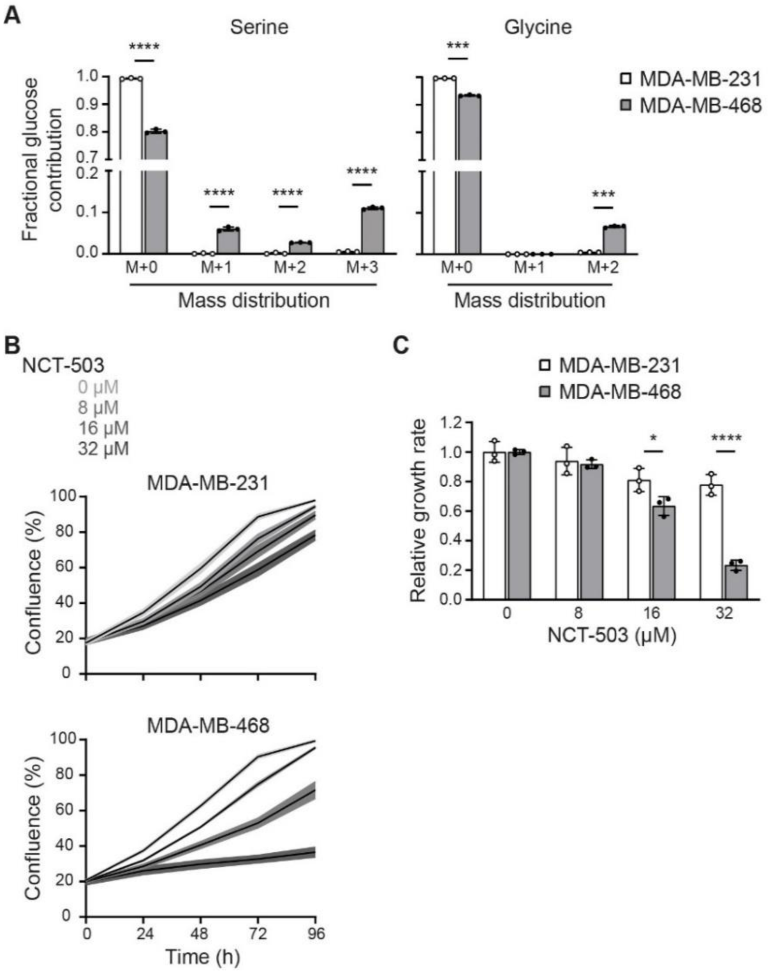


**Figure S2. MDA-MB-231 and MDA-MB-468 distinguish in being serine/glycine uptake versus serine/glycine synthesis dependent, respectively. (A)** Carbon incorporation from $^{13}C_6$-glucose into serine and glycine in MDA-MB-231, a serine/glycine uptake dependent BRCA cell line, and MDA-MB-468, a serine/glycine synthesis dependent BRCA cell line (n = 3, Multiple t-test). **(B)** Proliferation, as determined by real-time monitoring of cell confluence (%), of MDA-MB-231 (upper) and MDA-MB-468 (lower) cells upon treatment with indicated concentrations of NCT-503 for 96 h. 1 representative result of three biological replicates, containing each at least three technical replicates, is shown. **(C)** Quantification of (B). Data are presented as growth rate relative to the control treatment (n = 3, Two-way ANOVA, Sidak's multiple comparisons test). In **(A-C)** data are presented as mean ± SD. *p < 0.05, ***p < 0.001, ****p < 0.0001.

**Figure S3**

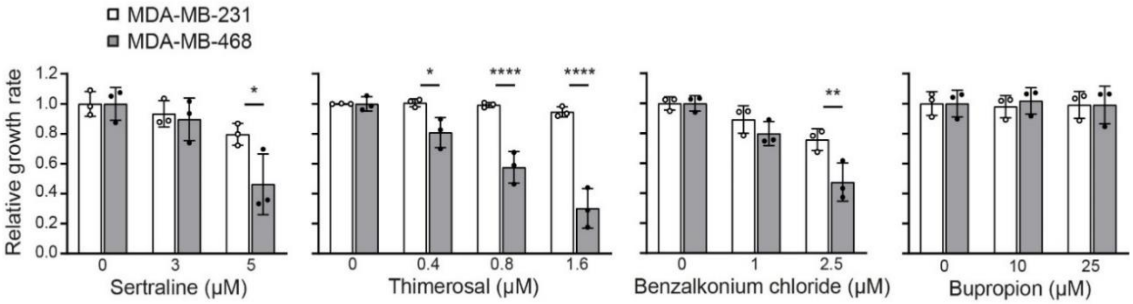


**Figure S3. Sertraline, thimerosal and benzalkonium chloride selectively decrease MDA-MB-468 cell proliferation.** Quantification of proliferation during 96 h, as determined by real-time monitoring of cell confluence (%), of MDA-MB-231 (white) and MDA-MB-468 (grey) cells upon treatment with indicated concentrations of sertraline, thimerosal, benzalkonium chloride and bupropion (left to right). Data are presented as growth rate relative to the control treatment (mean ± SD). *p < 0.05, **p < 0.01, ****p < 0.0001, Two-way ANOVA, Sidak's multiple comparisons test.

**Figure S4**

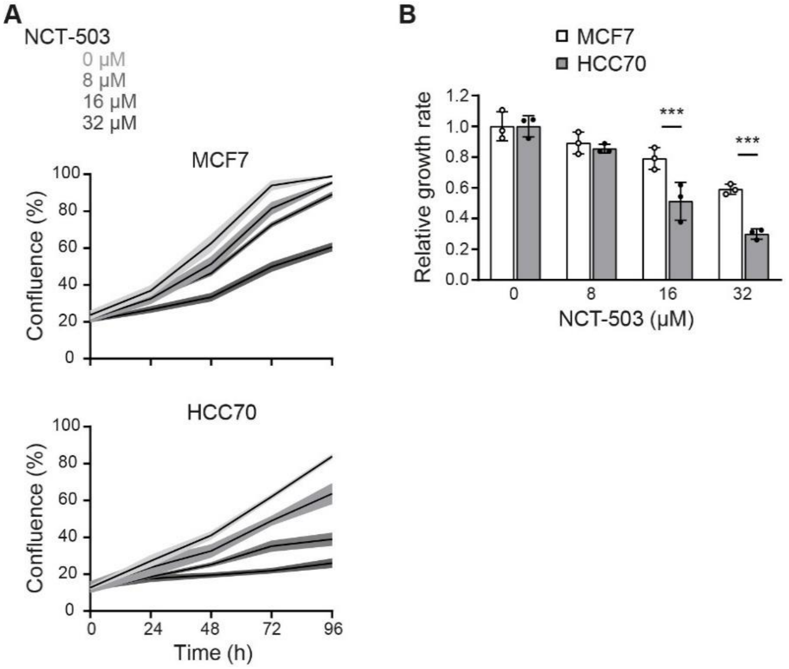


**Figure S4. NCT-503 inhibits proliferation of HCC70 more than of MCF7. (A)** Proliferation, as determined by real-time monitoring of cell confluence (%), of MCF7 (upper), a serine/glycine uptake dependent BRCA cell line, and HCC70 (lower), a serine/glycine synthesis dependent BRCA cell line, cells upon treatment with indicated concentrations of NCT-503 for 96 h. 1 representative result of three biological replicates, containing each at least three technical replicates, is shown. **(B)** Quantification of (A). Data are presented as growth rate relative to the control treatment ($n = 3$, Two-way ANOVA, Sidak's multiple comparisons test). In **(A-B)** data are presented as mean ± SD. ***$p < 0.001$.

#### Figure S5

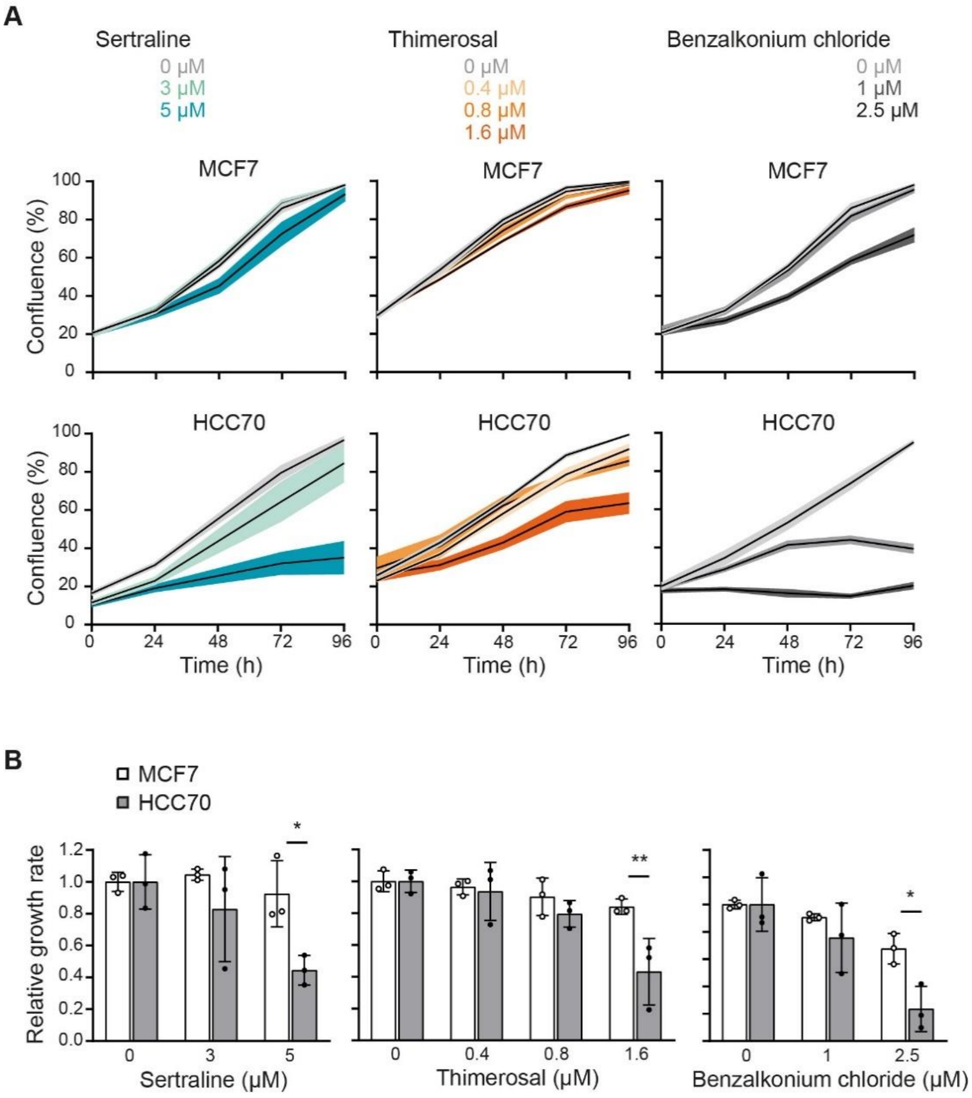


**Figure S5. Sertraline, thimerosal and benzalkonium chloride selectively impair HCC70 cell proliferation. (A)** Proliferation during 96 h, as determined by real-time monitoring of cell confluence (%), of MCF7 (upper) and HCC70 (lower) cells upon treatment with indicated concentrations of sertraline, thimerosal, benzalkonium chloride (left to right). **(B)** Quantification of (A). Data are presented as growth rate relative to the control treatment (n = 3, Two-way ANOVA, Sidak's multiple comparisons test). In **(A-B)** data are presented as mean ± SD. *$p < 0.05$, **$p < 0.01$.

Figure S6

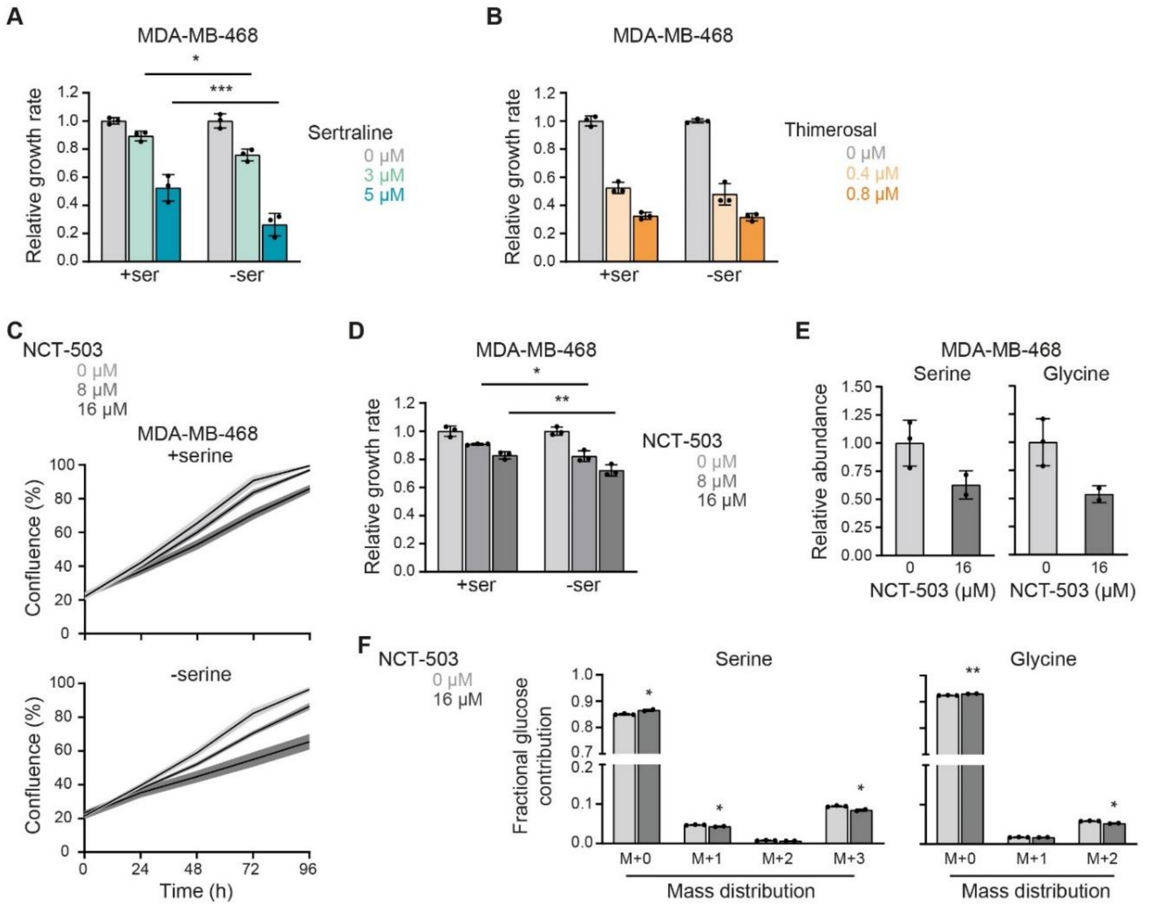


**Figure S6. Sertraline and thimerosal act similar as NCT-503, an already established inhibitor of serine/glycine synthesis. (A-B)** Quantification of proliferation during 96 h, as determined by real-time monitoring of cell confluence (%), of MDA-MB-468 cells cultured in DMEM with or without serine (400 μM) and treated with indicated concentrations of sertraline **(A)** or thimerosal **(B)**. Data are presented as growth rate relative to the control treatment (n = 3, Two-way ANOVA, Sidak's multiple comparisons test). **(C)** Proliferation during 96 h, as determined by real-time monitoring of cell confluence (%), of MDA-MB-468 cells cultured in DMEM with or without serine (400 μM) and treated with indicated concentrations of NCT-503. 1 representative result of three biological replicates, containing each at least three technical replicates, is shown. **(D)** Quantification of (C). Data are presented as growth rate relative to the control treatment (n = 3, Two-way ANOVA, Sidak's multiple comparisons test). **(E)** Relative abundance of intracellular serine and glycine in MDA-MB-468 cells treated with control (DMSO) or NCT-503 (16 μM) for 24 h (n = 2-3, Student's t-test). **(F)** Serine and glycine mass distribution showing the fractional glucose contribution of each mass upon treatment of MDA-MB-468 cells with control (DMSO) or NCT-503 (16 μM) (n = 2-3, Multiple t-test). In (A-F) data are presented as mean ± SD. *p < 0.05, **p < 0.01, ***p < 0.001.

**Figure S7**

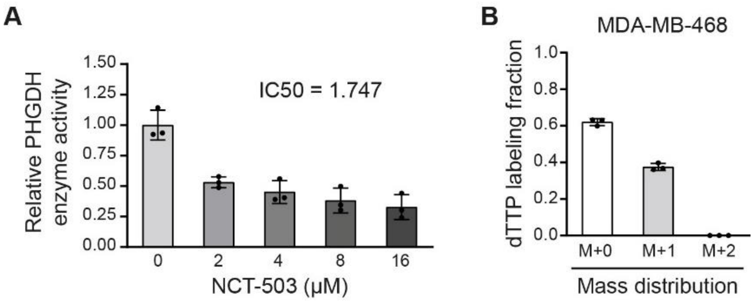


**Figure S7. Sertraline is an inhibitor of SHMT2 in MDA-MB-468 cells. (A)** PHGDH (= phosphoglycerate dehydrogenase) *in vitro* enzymatic assay, measuring PHGDH activity upon addition of indicated concentrations of NCT-503. Values are presented relative to the control (n = 3). **(B)** Deuterium label ($^2$H) incorporation into thymidine (dTTP) in MDA-MB-468 cells incubated with [2,3,3-$^2$H]-serine (deuterium labeled serine) for 48 h. Only M+1 dTTP (SHMT2), and no M+2 dTTP (SHMT1), was detected (n = 3). In **(A-B)** data are presented as mean ± SD.

**Figure S8**

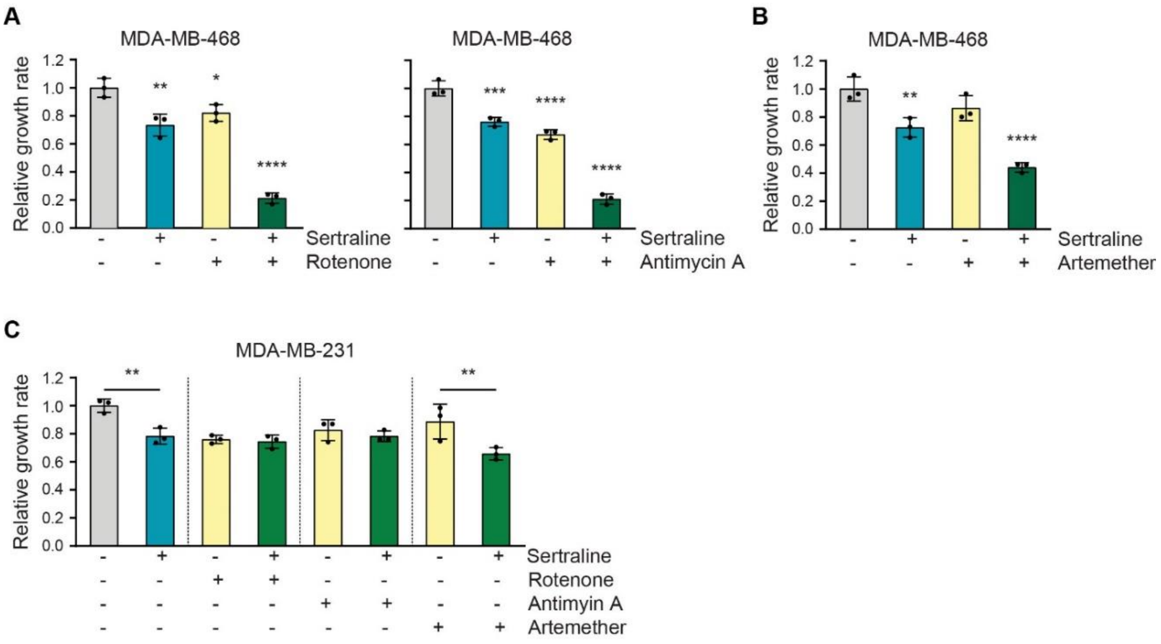


**Figure S8. Combining sertraline with drugs causing general mitochondrial dysfunction further decreases proliferation of MDA-MB-468 cells. (A-B)** Quantification of proliferation during 96 h, as determined by real-time monitoring of cell confluence (%), of MDA-MB-468 cells upon treatment with sertraline (5 µM) (blue) and/or (green/yellow) rotenone (50 nM) **(A)**, antimycin A (50 nM) (A) or artemether (80 µM) **(B)** (n = 3, Two-way ANOVA, Dunnet's multiple comparisons test). **(C)** Quantification of proliferation during 96 h, as determined by real-time monitoring of cell confluence (%), of MDA-MB-231 cells upon treatment with sertraline (5 µM) (blue) and/or (green/yellow) rotenone (50 nM), antimycin A (50 nM) or artemether (80 µM) (n = 3, One-way ANOVA, Sidak's multiple comparisons test). In **(A-C)** data are presented as growth rate relative to the control treatment (mean ± SD). *$p < 0.05$, **$p < 0.01$, ***$p < 0.001$, ****$p < 0.0001$.

Figure S9

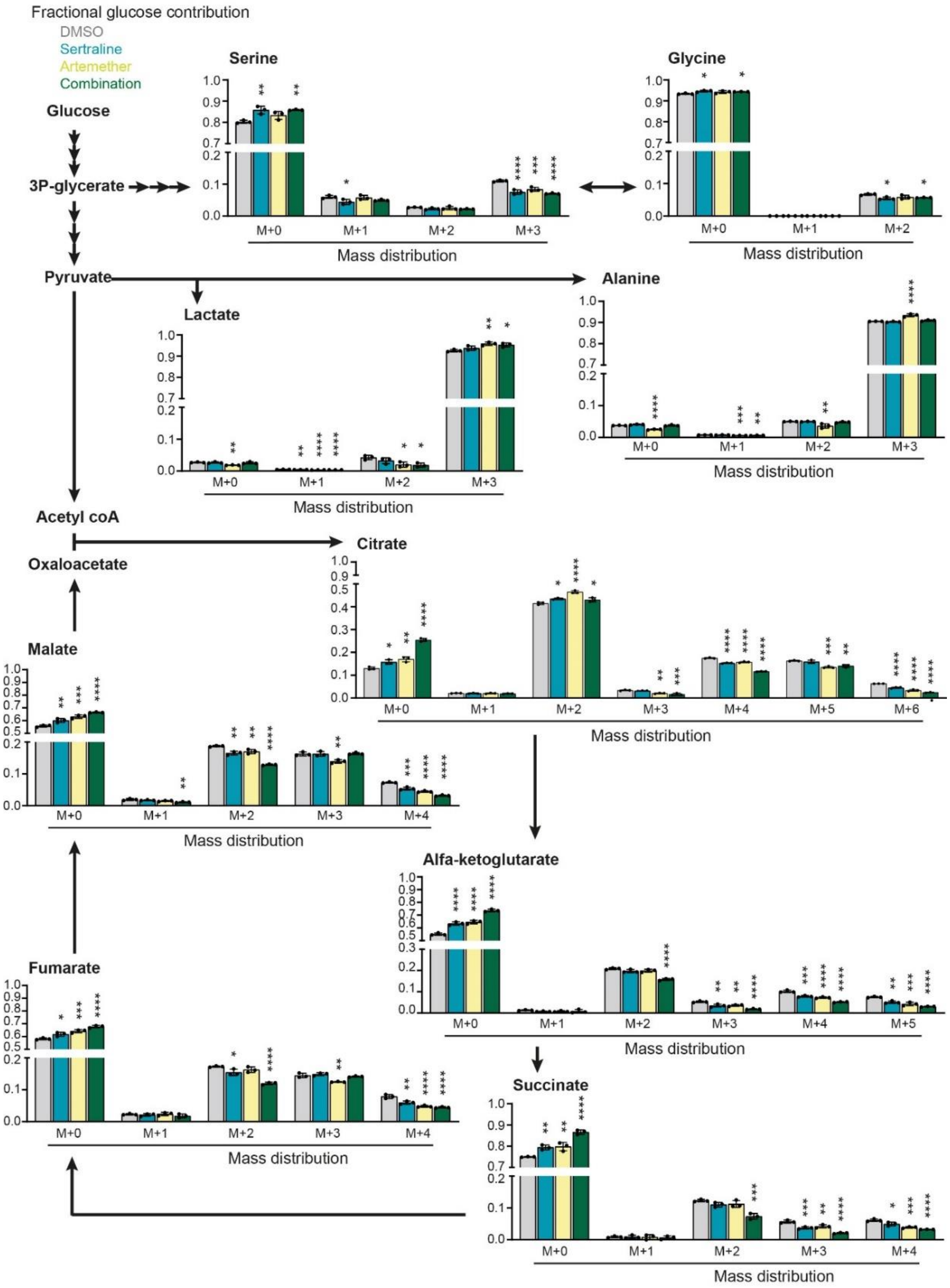


**Figure S9. The sertraline-artemether combination further metabolically disrupts cancer cells.** Carbon incorporation from $^{13}C_6$-glucose into downstream metabolites showing that both compounds work together in reducing the proliferation of MDA-MB-468 by decreasing both the amount of labeled TCA cycle metabolites and the flux through serine/glycine synthesis. The effect on serine/glycine synthesis is due to sertraline rather than artemether (blue = sertraline, yellow = artemether, green = combination) (n = 3, Two-way ANOVA, Dunnett's multiple comparisons test). Data are presented as mean ± SD. *$p<0.05$, **$p<0.01$, ***$p<0.001$, ****$p<0.0001$.

**Figure S10**

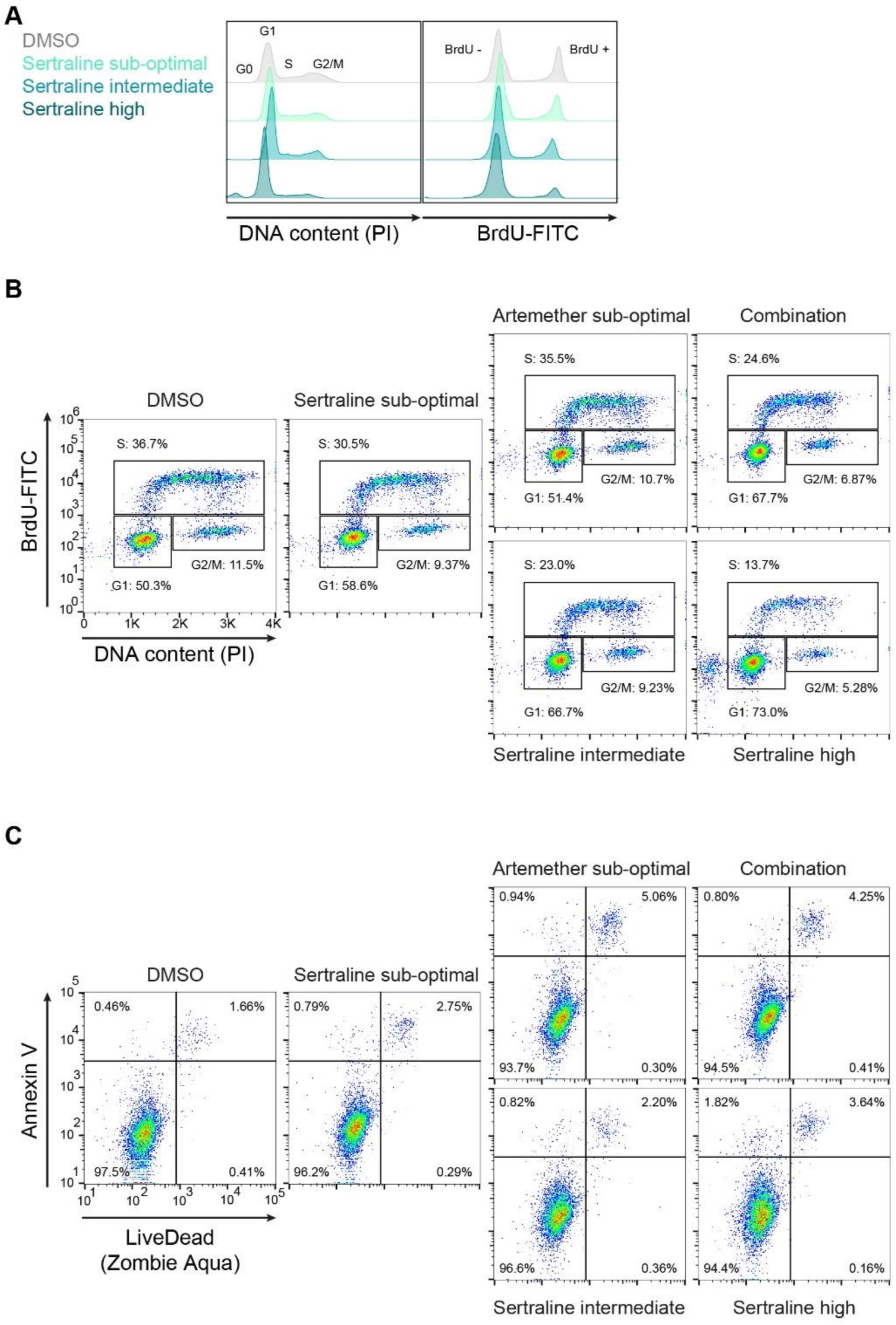


**Figure S10. The sertraline-artemether combination causes G1-S cell cycle arrest. (A)** Histograms showing PI cell cycle analysis (left) and BrdU incorporation (right) of MDA-MB-468 cells treated with DMSO or sertraline (sub-optimal = 5 µM; intermediate = 7.5 µM; high = 10 µM) for 24

h. 1 representative result of three biological replicates is shown. **(B)** Representative FACS dot-plot for BRDU-PI cell cycle analysis of MDA-MB-468 cells treated with DMSO, sertraline (sub-optimal = 5 µM; intermediate = 7.5 µM; high = 10 µM) and/or artemether (80 µM) for 24 h and incubated with BrdU for 1 h. Cells were gated on viable cells, including only the single cells. **(C)** Representative FACS dot-plot for Annexin V (PE) – Zombie Aqua (eFluor506) cell viability staining of MDA-MB-468 cells treated with DMSO, sertraline (sub-optimal = 5 µM; intermediate = 7.5 µM; high = 10 µM) and/or artemether (80 µM) for 24 h. Cells were gated on viable cells, including only the single cells.
